# Systematic mapping of MCU-mediated mitochondrial calcium signaling networks

**DOI:** 10.1101/2024.02.20.581153

**Authors:** Hilda Delgado de la Herran, Denis Vecellio Reane, Yiming Cheng, Máté Katona, Fabian Hosp, Elisa Greotti, Jennifer Wettmarshausen, Maria Patron, Hermine Mohr, Natalia Prudente de Mello, Margarita Chudenkova, Matteo Gorza, Safal Walia, Michael Sheng-Fu Feng, Anja Leimpek, Dirk Mielenz, Natalia S. Pellegata, Thomas Langer, György Hajnóczky, Matthias Mann, Marta Murgia, Fabiana Perocchi

**Affiliations:** Institute for Diabetes and Obesity, Helmholtz Diabetes Center, Helmholtz Zentrum Munich, Munich, Germany; Department of Pathology, Anatomy, and Cell Biology, MitoCare Center, Thomas Jefferson University, Philadelphia, USA; Department of Proteomics and Signal Transduction, Max Planck Institute of Biochemistry, Martinsried, Germany; Neuroscience Institute, National Research Council of Italy, Padua, Italy; Department of Biomedical Sciences, University of Padova, Padua, Italy; Padova Neuroscience Center, University of Padova, Padua, Italy; Institute for Genetics, Cologne Excellence Cluster on Cellular Stress Responses in Aging-Associated Diseases, Center for Molecular Medicine, University of Cologne, Cologne, Germany; Max Planck Institute for Biology of Aging, Cologne, Germany; Institute of Diabetes and Cancer, Helmholtz Center Munich, Munich, Germany; Division of Molecular Immunology, University of Erlangen, Nikolaus-Fiebiger-Zentrum, FAU Erlangen-Nürnberg, Erlangen, Germany; Department of Biology and Biotechnology, University of Pavia, Pavia, Italy; Faculty of Health Sciences, Novo Nordisk Foundation Center for Protein Research, University of Copenhagen, Copenhagen, Denmark; Institute of Neuronal Cell Biology, Technical University of Munich, Munich, Germany; Munich Cluster for Systems Neurology, Munich, Germany

**Keywords:** Calcium Signaling, Mitochondria, Mitochondrial Calcium Uniporter, Organelle, Proteomics

## Abstract

The Mitochondrial Ca^2+^ Uniporter Channel (MCUC) allows calcium entry into the mitochondrial matrix to regulate energy metabolism but also cell death. Although, several MCUC components have been identified, the molecular basis of mitochondrial Ca^2+^ signaling networks and their remodeling upon changes in uniporter activity have not been systematically assessed. Using an unbiased and quantitative proteomic approach, we map the MCUC interactome in HEK293 cells under physiological conditions and upon chronic loss or gain of mitochondrial Ca^2+^ uptake. Besides all previously known subunits of the uniporter, we identify 89 high-confidence interactors linking MCUC to several mitochondrial complexes and pathways, half of which are currently linked to metabolic, neurological, and immunological diseases. As a proof-of-concept, we validate EFHD1 as a binding partner of MCU, EMRE and MCUB with a MICU1-dependent inhibitory effect on Ca^2+^ uptake. To investigate compensatory mechanisms and functional consequences of mitochondrial Ca^2+^ dyshomeostasis, we systematically survey the MCU interactome upon silencing of EMRE, MCUB, MICU1 or MICU2. We observe profound changes in the MCU interconnectivity, whereby downregulation of EMRE reduces the number of MCU interactors of over 10-fold, while silencing of MCUB leads to a wider functional network linking MCU to mitochondrial stress response pathways and cell death. Altogether our study provides a comprehensive map of MCUC protein-protein interactions and a rich, high-confidence resource that can be explored to gain insights into the players and mechanisms involved in calcium signal transduction cascades and their relevance in human diseases.

## INTRODUCTION

For decades, mitochondria have been recognized as key players in Ca^2+^-mediated signal transduction cascades, decoding the spatio-temporal dynamics of intracellular Ca^2+^ signals (Hajnóczky *et al*, 1995; Spät *et al*, 2008; Kaftan *et al*, 2000). This property enables the organelle to regulate the metabolic state of the cell, its growth, fate, and overall survival (Rizzuto *et al*, 2012). Indeed, changes in cytosolic Ca^2+^ concentration ([Ca^2+^]_cyt_) are promptly transferred into the mitochondrial matrix through an electrogenic pathway powered primarily by the mitochondrial membrane potential and mediated by MCUC, a highly selective calcium channel located at the inner mitochondrial membrane (IMM) (Kirichok *et al*, 2004; DeLuca & Engstrom, 1961; Vasington & Murphy, 1962). A transient elevation of matrix Ca^2+^ concentration ([Ca^2+^]_mt_) is then efficiently coupled to the regulation of mitochondrial bioenergetics to match the metabolic demands of cells and tissues (Tsai *et al*, 2022; Fecher *et al*, 2019; Fieni *et al*, 2012; Paillard *et al*, 2017). However, when sustained, mitochondrial Ca^2+^ (mt-Ca^2+^) uptake can trigger oxidative stress and mitochondrial swelling, with consequent activation of cell death pathways (Rizzuto *et al*, 2012). Accordingly, dyshomeostasis of mt-Ca^2+^ has already been implicated in numerous diseases including diabetes, neurodegeneration, stroke, heart failure, inflammation, muscular atrophy, and cancer (Garbincius & Elrod, 2022). Therefore, elucidating the molecular determinants of channel activity and MCUC-dependent control of mitochondrial functions represents an important milestone in cell physiology and pathophysiology.

Unbiased computational and experimental analyses that leveraged co-evolution, co-expression, and organellar proteomics led to the discovery of MICU1 (<u>M</u>itochondrial <u>C</u>alcium <u>U</u>ptake <u>1</u>) (Perocchi *et al*, 2010) and MCU (<u>M</u>itochondrial <u>C</u>alcium <u>U</u>niporter) (De Stefani *et al*, 2011; Baughman *et al*, 2011), as Ca^2+^-dependent regulator and pore-forming subunit of the uniporter, respectively. These findings paved the way for the characterization of additional binding partners, including the MCU-dominant negative beta subunit (MCUB)(Raffaello *et al*, 2013), a family of MICU1 paralogs and splice variants (MICUs) (Plovanich *et al*, 2013; Patron *et al*, 2019; Vecellio Reane *et al*, 2016; Patron *et al*, 2014), and EMRE (<u>E</u>ssential <u>M</u>CU <u>Re</u>gulator)(Sancak *et al*, 2013). However, although several channel forming elements are currently known, MCUC-dependent Ca^2+^ signaling networks in mitochondria have not yet been assessed. A few studies have applied affinity purification (AP) coupled with quantitative mass spectrometry (MS) for an unbiased mapping of MCU protein-protein interactions (PPIs) (Sancak *et al*, 2013; Austin *et al*, 2022; Antonicka *et al*, 2020) but failed to identify functional associations with other mitochondrial proteins besides the known MCUC subunits. Instead, recent evidence indicates that MCUC is part of a broader functional network and can dynamically interact with other mitochondrial complexes, for example with members of the respiratory chain complex I (RCCI) (Balderas *et al*, 2022), the ATP-dependent proteolytic complex (König *et al*, 2016; Tsai *et al*, 2017), and the mitochondrial contact site and cristae organization system (MICOS) (Tomar *et al*, 2023; Gottschalk *et al*, 2019). In this way, the organelle can regulate the stability, assembly and activity of the uniporter, which in turn influence mitochondrial membrane structure and energy synthesis, and compensates for mitochondrial dysfunctions. Nevertheless, a systematic and comprehensive analysis of the MCUC protein interaction landscape and signal transduction networks, both under resting conditions and perturbed mt-Ca^2+^ homeostasis, is still lacking.

Here, we devised a biochemical strategy to characterize the MCUC interactome in human cells with high confidence and resolution under physiological conditions and following genetic perturbations. Tandem affinity purifications (TAPs) coupled with quantitative and integrative liquid chromatography-tandem mass spectrometry (LC-MS/MS) analyses led us to map 139 statistically significant PPIs between 95 mitochondrial proteins in HEK293, including all currently known members of the uniporter complex. We were able to capture all previously described interactions between MCUC, RCCI, MICOS, and several mitochondrial proteases, as well as novel molecular links between mt-Ca^2+^ signaling, organelle dynamics, biogenesis, and apoptosis. Among the MCUC binding partners, we also identified a handful of hitherto poorly characterized proteins, for example the EF-hand domain-containing protein D1 (EFHD1, also known as Swiprosin-2 or Mitocalcin) (Hou *et al*, 2016; Dütting *et al*, 2011). We corroborate a role for EFHD1 as an inhibitor of MCU-mediated mt-Ca^2+^ uptake, a function that appears to be dependent on the presence of MICU1. Next, to investigate the molecular basis of compensatory mechanisms and functional consequences of MCUC remodeling upon loss-of-function (LOF) of its components, we performed LC-MS/MS analyses of TAPs from MCU-tagged HEK293 cells upon silencing of EMRE, MCUB, MICU1 or MICU2. Upon EMRE knockdown (KD) the number of PPIs was dramatically reduced, possibly due to the inability of MCU to form active channels. In contrast, MCUB KD, which increases mt-Ca^2+^ uptake, led to an expansion and greater interconnection of the MCU protein network and prevented MCU from forming high molecular weight (MW) macromolecular complexes, in both human cells and mouse tissues. Altogether, our dataset represents a rich, high-confidence resource that can be explored to discover novel molecular links between MCUC-mediated Ca^2+^ entry and organelle biology, as well as providing new insights into unexplored mt-Ca^2+^ signaling and disease associations.

## RESULTS

### Systematic and unbiased identification of MCUC protein-protein interactions

We devised a biochemical workflow to achieve an unbiased and comprehensive characterization of MCUC-specific PPIs in human cells under near-physiological conditions (**Fig. 1**). To this goal, MCU, MCUB or EMRE proteins were fused at the C-terminus to a StrepII-HA-His_6_ tag and stably integrated into Flp-In T-REx HEK293 cells (parental line) under the control of a tetracycline-inducible promoter. This allowed the targeted integration of each bait protein at a single transcriptionally active genomic locus that is the same in every cell line and ensures a near-physiological expression level of the bait proteins. The latter is essential to minimize false positive associations and accurately define the endogenous protein composition and interaction network of MCUC. By encompassing all three membrane-spanning subunits of MCUC as baits, we aimed to increase the coverage of copurifying proteins across all mitochondrial sub-compartments. As shown in **Fig. 2A** and **Fig. S1**, we confirmed that tetracycline-dependent induction of baits did not affect the endogenous level of other MCUC subunits, except for EMRE. Upon expression of tagged MCU and EMRE, the level of endogenous EMRE was increased and decreased, respectively, which is consistent with previous findings indicating a tight regulation of its stability by MCU (Sancak *et al*, 2013; Tsai *et al*, 2017; König *et al*, 2016).

**Figure 1.**
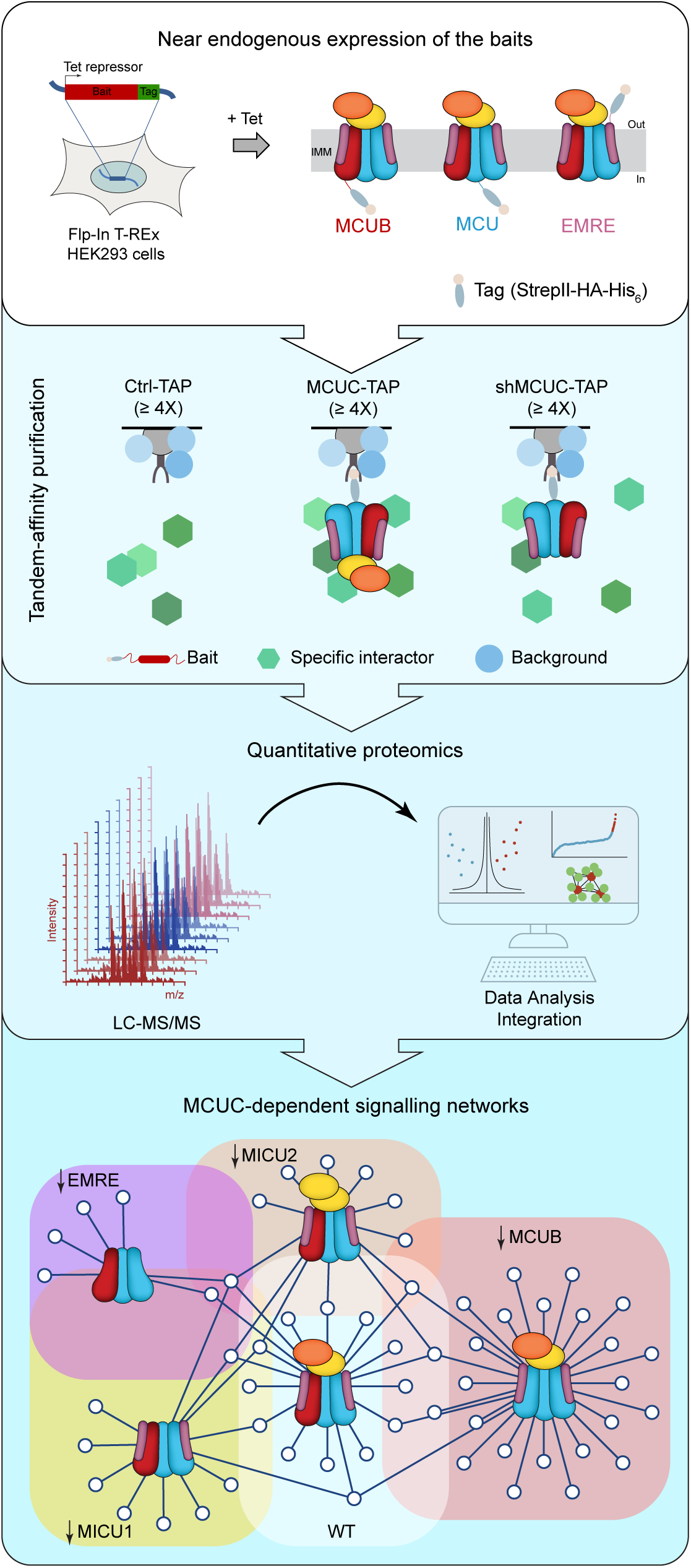
Proteomics approach to characterize the MCUC interactome. Tandem affinity purifications (TAPs) from Flp-In T-REx HEK293 cells that are wild-type (Ctrl) or expressing MCU, MCUB, and EMRE as baits. *Tet*, tetracyclin; *IMM*, inner mitochondrial membrane; *StrepII*, streptavidin II tag; *His*_6_, polyhistidine tag; *HA*, hemagglutinin tag; *LC-MS/MS*, liquid chromatography-tandem mass spectrometry; *[Ca^2+^]*_mt_, mitochondrial Ca^2+^ concentration.

**Figure 2.**
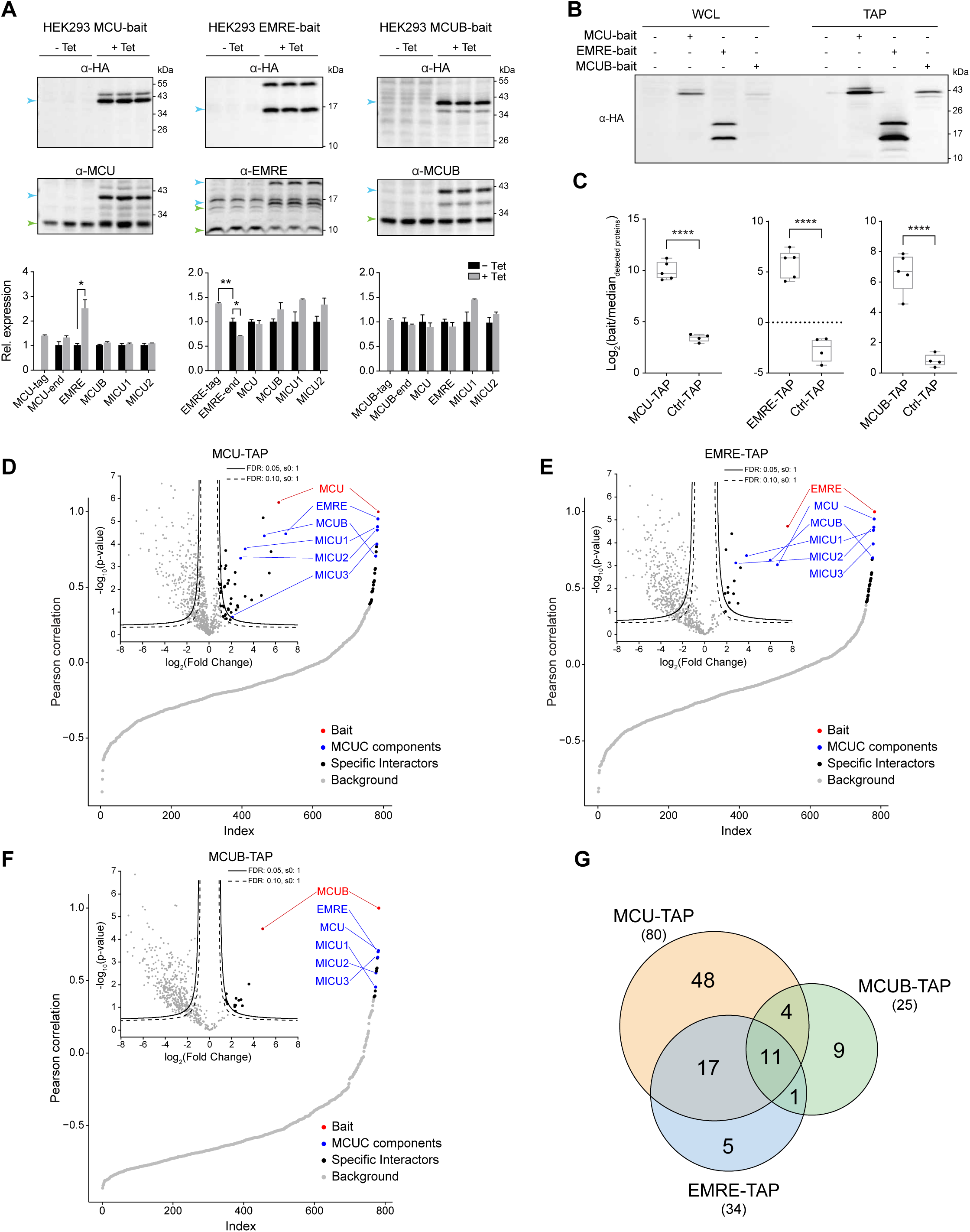
Unbiased and systematic identification of endogenous MCU, MCUB, and EMRE protein interactors. **A**, Protein expression of known MCUC subunits upon bait induction. *Upper*, immunoblot analysis of whole cell lysates before (–Tet) and after (+Tet) tetracycline-driven expression of MCU, EMRE, and MCUB. Green and blue arrows refer to endogenous and exogenous protein levels, respectively. *Lower*, Quantification of MCUC protein expression (–tag and –end refer to exogenous and endogenous protein level, respectively) normalized to ACTIN (loading control) and relative to –Tet condition (mean ± SEM; n= 3); student’s t-test; *p< 0.05, **p< 0.01. **B**, Bait enrichment after TAP from whole cell lysates (WCL). **C**, Intensity-based absolute quantification of baits over the median intensity of all detected proteins in biological replicates of TAPs (mean ± SEM; n≥ 4); student’s t-test; ****p< 0.0001. **D**, **E**, **F**, Global Pearson’s correlation rank and Hawaii plots of mitochondrial proteins enriched in MCU-bait (D), EMRE-bait (E), and MCUB-bait (F) TAPs. Continuous and dashed lines indicate specific interaction partners defined by permutation-based FDR thresholds of 0.05 or 0.10, respectively. **G**, Overlap of MCUC interactors from all three baits. See also **Figure S1** and **Table S1**.

We employed a TAP strategy based on Strep and poly-histidine tags to enrich for interacting protein partners of MCUC. Immunoblot analyses validated that each bait could be efficiently purified from whole cell lysates (**Fig. 2B**). For a systematic and unbiased identification and quantification of proteins that co-precipitated with each bait, we performed LC-MS/MS analysis with at least four biological replicates. Importantly, we included TAPs from the parental line, hereby used as negative control (Ctrl-TAP) to discriminate specific interactions from spurious ones. As a result, we found that the relative abundance of MCU, MCUB, and EMRE was consistently and significantly higher in TAPs from bait-expressing cells than in control samples, when compared to the median intensity of all quantified proteins (**Fig. 2C**). To define significant and specific MCUC interactors we filtered our dataset for mitochondrial proteins and ran a quantitative bait-prey co-enrichment analysis based on a two-tailed Welch’s t-test, a within-group variance (s0) of 1, and a permutation-based false discovery rate (FDR) of either 0.05 (“high-confidence”) or 0.10 (“medium-confidence”) (**Fig. 2D-F**). In addition, Hein et al. have previously demonstrated that the intensity profiles of interacting proteins are correlated (Hein *et al*, 2015). Therefore, as additional classifiers to identify MCUC interactors, we performed “local” and “global” correlation analyses, comparing the similarity of protein intensity profiles across either pairs of bait and control TAPs, or across all measured samples (Ctrl-TAPs, MCU-TAPs, MCUB-TAPs, and EMRE-TAPs), respectively.

With the term “interactor” or “prey” we refer to any protein that stably or dynamically binds MCUC directly or indirectly. Indirect binding can occur when baits and preys are in close proximity without engaging in direct physical interactions. These associations might be facilitated by subunits of a protein complex or members of a pathway that directly interact with MCUC. Examples include connections between mitochondrial protease complexes and MCUC (König *et al*, 2016; Tsai *et al*, 2017, 2022) or between MCU and RCCI (Balderas *et al*, 2022). The joint analysis of all datasets yielded a total of 139 interactions among 95 mitochondrial proteins and successfully recovered all currently known subunits of the MCUC, namely MCU, EMRE, MCUB, MICU1, MICU2, and MICU3 (**Fig. 2G** and **Table S1**). Among the MCUC interactors, 33 were found to be shared between at least two baits, and 11 were common to all three conditions. Interestingly, more than 50% of all preys were bait-specific, suggesting their involvement in the selective maturation and regulation of MCU, EMRE, or MCUB.

### The MCUC protein network identifies molecular links between Ca^2+^ and mitochondrial functions

To globally analyze MCUC-specific PPIs, we assessed the biochemical, functional, and evolutionary properties of all identified protein partners (**Fig. 3** and **Table S1-3**). Roughly 50% of our interactome consisted of proteins with at least one predicted transmembrane domain, for example subunits of the outer (TOM) and inner (TIM) membrane protein translocases and the oxidative phosphorylation (OXPHOS) machinery. The other half were soluble proteins including components of the MICUs family, the tricarboxylic acid cycle (TCA cycle), and the mitochondrial DNA (mt-DNA) maintenance and expression system (**Fig. 3A**). Accordingly, the uniporter engaged in interactions with proteins localized in all four submitochondrial compartments (**Fig. 3B**). MCU and EMRE, which represent the minimal functional unit of MCUC (Kovács-Bogdán *et al*, 2014; Pittis *et al*, 2020), sampled largely similar environments although exposing their tags on opposite sides of the IMM. Interestingly, they also mediated functional associations with proteins located at the outer mitochondrial membrane (OMM) and involved in mitochondrial morphology, apoptosis, and dynamics, such as FIS1, BNIP3L, and PARK7. Besides subunits of MCUC, the identified interactors encompassed a wide spectrum of mitochondrial functions, with several not yet known to be linked to Ca^2+^ signaling (**Fig. 3C**).

**Figure 3.**
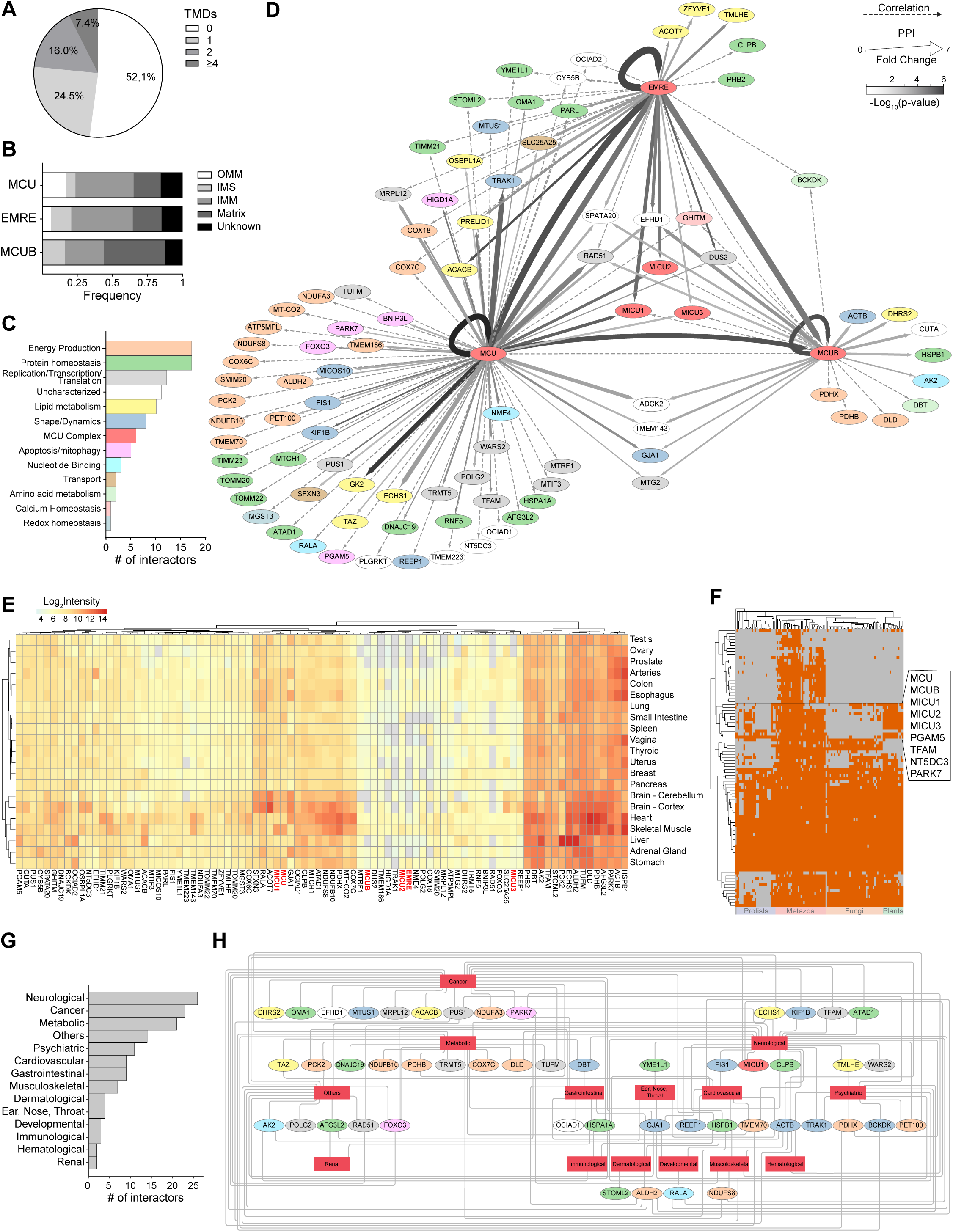
Systems-wide analysis of the MCUC protein network. **A**, Percentage of MCUC interactors with transmembrane domains (TMDs). **B**, Distribution of bait-specific interactors across sub-mitochondrial compartments. *OMM*, outer mitochondrial membrane; *IMS*, intermembrane space; *IMM*, inner mitochondrial membrane; *Unknown*, missing information or multi-localized proteins. **C**, Distribution of MCUC interactors into functional categories. **D**, Network of MCUC protein-protein-interactions (PPIs) defined by quantitative co-enrichment (solid line), local and global correlation analyses (dashed line). Color and thickness of solid lines indicate statistical significance (p-value) and enrichment (fold change), respectively, whereas the color of each node refers to the protein functional annotation. **E**, Heatmap of relative protein expression for MCUC interactors in human tissues. **F**, Phylogenetic profiles of MCUC interactors across 120 eukaryotic organisms and *E. coli.* **G**, Absolute frequency histogram for known disease classes associated to MCUC interactors. **H**, Gene-disease network analysis of MCUC interactors. See also **Table S1**, **Table S2**, and **Table S3.**

We then mapped the MCUC interactome as a protein network, whereby each node corresponds to a prey connected to a given bait through either evidence of co-enrichment or profile correlations (**Fig. 3D**). Reassuringly, each bait significantly interacted with itself and with all known MCUC members. To further corroborate the high quality and coverage of our dataset, we first searched the literature for experimental evidence of physical and functional associations between the uniporter and mitochondrial proteins in our network. Among shared interactors we identified GHITM, also known as TMBIM5 or MICS1. This protein was recently characterized as a Ca^2+^/H^+^ exchanger of the IMM (Zhang *et al*, 2022), able to shape mt-Ca^2+^ cycling by inhibiting the activity of the m-AAA protease AFG3L2 (Austin *et al*, 2022; Patron *et al*, 2022). We also identified AFG3L2 and the YME1L proteolytic hub (YME1L1, PARL, and STOML2) (Wai *et al*, 2016), all previously implicated in MCUC processing, assembling and degradation (König *et al*, 2016; Tsai *et al*, 2017, 2022). To test the usefulness of the MCUC protein network as a resource for the identification of novel research directions and regulatory mechanisms, we also mined our dataset for associations between the uniporter and mitochondrial functions that are known to be controlled by Ca^2+^ but for which the molecular players remain poorly characterized. For instance, the MCUC interactome included several proteins (FIS1, KIF1B, GJA1, and MTUS1) involved in mitochondrial ultrastructural organization, shape and dynamics. This supports the notion that mt-Ca^2+^ signaling can affect the organelle’s morphology through the regulation of mitochondrial fusion and fission (Zhao *et al*, 2015). In addition, our results raise the hypothesis that MCU, and thus mt-Ca^2+^ signaling, could regulate contact sites between inner and outer mitochondrial membranes through interactions with members of the MICOS complex such as MICOS10 (Rampelt *et al*, 2022). Notably, MICU1 was recently found to bind the MICOS complex and regulate mitochondrial membrane dynamics independently of matrix Ca^2+^ uptake, while MCU was shown to re-localize at the cristae junctions upon an increase of [Ca^2+^] in the IMS (Tomar *et al*, 2023; Gottschalk *et al*, 2019). Given mt-Ca^2+^ signaling is a central regulator of oxidative metabolism and ATP production (Hajnóczky *et al*, 1995; Jouaville *et al*, 1999), we also looked for molecular players coupling MCUC with mitochondrial bioenergetics. Remarkably, we found that the MCUC network was especially enriched in PPIs with components of RCCI (e.g. NDUFA3, NDUFB10, NDUFS8), RCCIV (e.g. COX6C, COX7C, COX18) and the TCA cycle (PDHB and PDHX). Supporting our findings, the RCCI assembly subunit NDUFA3 was recently shown to bind MCU, an interaction between complexes that regulates uniporter level and activity to maintain bioenergetic homeostasis (Balderas *et al*, 2022). To the same goal, during Ca^2+^ signaling the activation of SLC25A25 allows matching cellular energy demand and supply by regulating the matrix adenyl nucleotide pool (del Arco *et al*, 2016). However, no previous evidence of an interaction between the uniporter and SLC25A25 has been reported. Instead, we found SLC25A25 to be significantly co-enriched with the MCU and EMRE baits possibly allowing them to co-localize in the same Ca^2+^ microdomain and co-activate in response to rise in Ca^2+^ concentration ([Ca^2+^]).

Next, we surveyed properties of mt-Ca^2+^ signaling by analyzing expression and evolutionary profiles of all MCUC interactors, as well as their involvement in human diseases (**Fig. 3E-H** and **Table S2, S3**). First, we quantified the relative protein level of each MCUC binding partner across healthy human tissues using the GTEx (Genotype-Tissue Expression) database (Jiang *et al*, 2020). Interestingly, we observed a highly heterogeneous tissue distribution of MCUC interactors, which is consistent with the wide range of biological processes linked to mt-Ca^2+^ in our network (**Fig. 3E**). These proteins encompass a wide range of functions, from fundamental mitochondrial processes like OXPHOS and mt-DNA replication, to more specialized roles, such as regulation of organelle shape and dynamics, as well as activation of cell death pathways. As expected, based on their role as the minimal unit of the uniporter, MCU and MICU1 showed a ubiquitous expression compared to tissue-specific regulators such as MICU2, MICU3 and MCUB. EMRE was often undetected due to its small size and highly hydrophobic nature, which makes it difficult to detect by MS analyses of whole proteomes. Next, we assessed the evolutionary conservation of each interactor across eukaryotes by mapping orthology relationships across 120 eukaryotic species inferred from Protphylo (Cheng & Perocchi, 2015). Comparative genomics analyses have been instrumental in identifying MCUC components (Perocchi *et al*, 2010; Baughman *et al*, 2011; De Stefani *et al*, 2011), which are present in all major eukaryotic groups, but mostly absent in protists and fungi (Pittis *et al*, 2020). However, while coevolution can be used to predict functional associations, functionally related proteins do not necessarily co-evolve, calling for complementary approaches, like ours, to reach a comprehensive map of mt-Ca^2+^ signaling networks. Namely, most of the proteins in our network did not exhibit similar phylogenetic profiles to MCU (**Fig. 3F**). Overall, they showed a patchy evolutionary conservation, with mitochondrial functional modules of ancient origin being highly conserved, while others present only in higher vertebrates. The former group could represent processes that were placed under the control of mt-Ca^2+^ early in evolution; the latter would possibly spotlight species-specific functions linked to Ca^2+^ signaling after the acquisition of MCU. Finally, we found that half of the MCUC interactors were linked to over 300 different disease phenotypes, spanning from neurological and metabolic disorders to cancer (**Fig. 3G**), including disorders related to mt-DNA homeostasis, whose pathophysiology has not yet been connected to dysfunctions in mt-Ca^2+^ signaling (**Fig. 3H**). Altogether, our analysis provides a high-confidence and comprehensive dataset of endogenous MCU, MCUB, and EMRE protein binding partners.

### Validation of newly identified MCUC interactors

To corroborate the quality of our resource, we experimentally validated a subset of specific interactions with different degrees of co-enrichment and correlation to MCU, EMRE, and MCUB baits. MICU3, which was not previously identified as a *bona fide* binding partner of MCUC in HEK293 cells (Sancak *et al*, 2013; Antonicka *et al*, 2020; Austin *et al*, 2022), resulted as a high-confidence interactor of all three baits in our dataset. Indeed, quantitative proteomic analyses of mitochondria have confirmed that MICU3 is expressed in HEK293 cells (Morgenstern *et al*, 2021), albeit in low amount when compared to other MCUC components (**Fig. 4A**). MICU3 is a mitochondrial protein located in the IMS and loosely attached to the IMM, which can form cysteine-mediated disulfide bonds with MICU1 and acts as an enhancer of MCUC thanks to its ability to sense [Ca^2+^] (**Fig. S2A-I**) (Patron *et al*, 2019). To assess whether MICU3 also played a functional role in the regulation of mt-Ca^2+^ uptake in HEK293 cells, we stably expressed either control (pLKO) or shRNAs targeting different regions of MICU3 mRNA (**Fig. 4B**). Notably, mitochondria of digitonin-permeabilized sh-MICU3 cells showed reduced Ca^2+^ uptake capacity compared to pLKO upon consecutive addition of exogenous Ca^2+^ (**Fig. 4C**). The Ca^2+^ clearance phenotype was strongly correlated to MICU3 protein level and therefore not likely due to an off-target effect. Next, we investigated the link between MCU, Tafazzin (TAZ), and the lipid transfer protein PRELI domain-containing protein 1 (PRELID1). Both proteins participate in the biosynthesis of cardiolipin (CL) (Tatsuta & Langer, 2017), which is required for the assembly and stability of numerous IMM protein complexes, including MCUC. Whereas cardiolipin metabolism was shown to affect MCU-dependent Ca^2+^ uptake (Gottschalk *et al*, 2019; Ghosh *et al*, 2020), the functional interactions involved have not been reported yet. Notably, we observed that PRELID1 KD resulted in a dramatic reduction of MCU protein level at both monomeric (**Fig. S2J**) and oligomeric states (**Fig. 4D**), confirming its role in the regulation of MCU stability. As a consequence of impaired MCUC assembly, histamine-stimulated si-PRELID1 HeLa cells showed a marked decrease in [Ca^2+^]_mt_ (**Fig. 4E**), without any obvious effect on [Ca^2+^]_cyt_ transients (**Fig. S2K**) and mitochondrial membrane potential (**Fig. S2L**).

**Figure 4.**
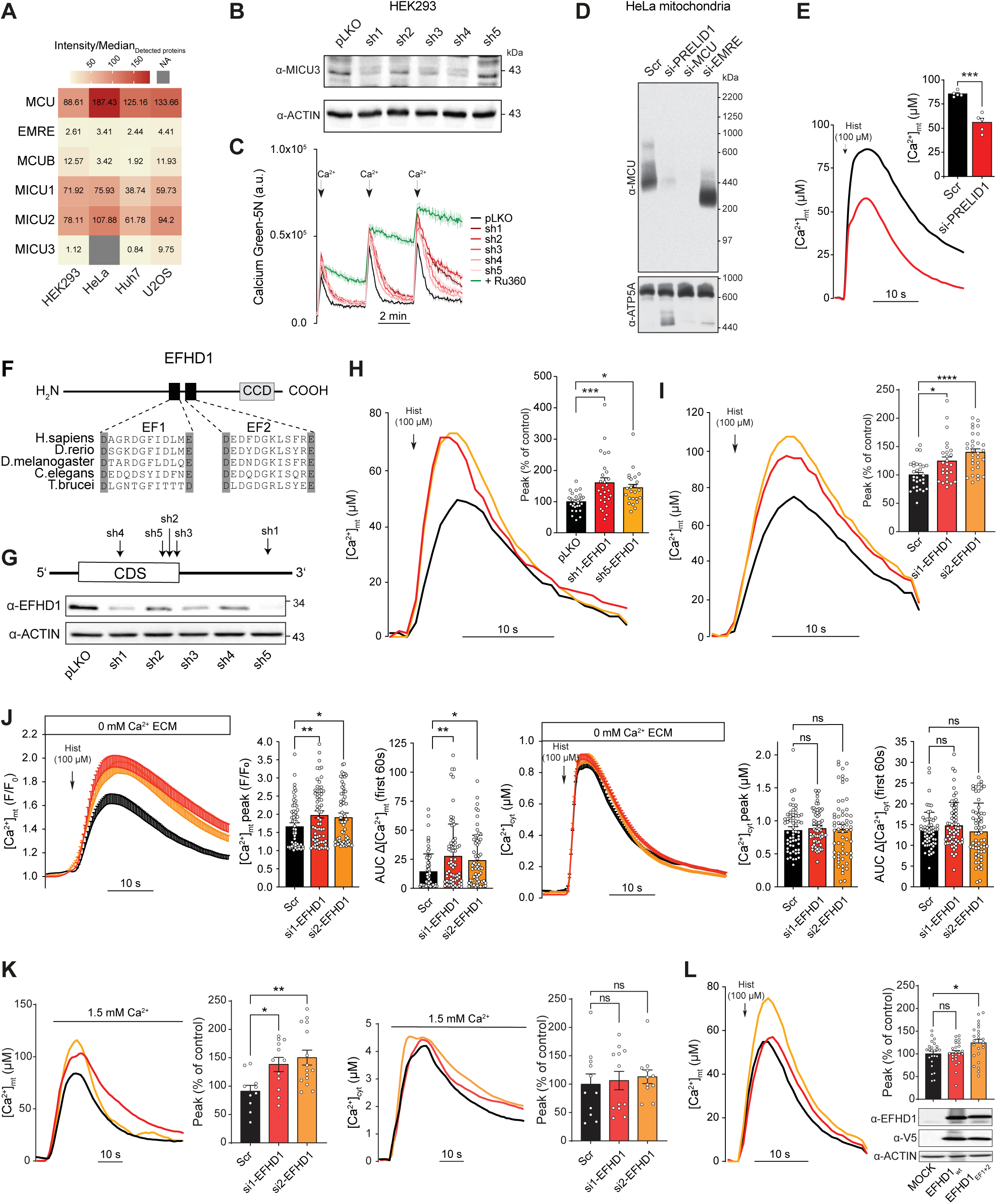
MICU3, PRELID1, and EFHD1 loss-of-function affect mt-Ca^2+^ homeostasis. **A**, MS-based quantification of MCUC components in mitochondria isolated from HEK293, HeLa, Huh7 and U2OS human cell lines as in MitoCoP (Morgenstern *et al*, 2021). *NA*, not assigned. **B**, Immunoblot analysis of MICU3 and ACTIN (loading control) protein level in whole cell lysates from HEK293 cells stably expressing either sh-MICU3 RNAs (sh1-5) or an empty vector (pLKO). **C**, Average kinetics of Ca^2+^ clearance by mitochondria of digitonin-permeabilized sh-MICU3 and pLKO HEK293 cells (arrow, injection of 40 μM CaCl_2_). Ru360 (10 µM) is used as a positive control for MCU-dependent Ca^2+^ uptake inhibition (mean ± SEM; n= 4). **D**, BN-PAGE analysis of mitochondria isolated from HeLa cells transfected with siRNAs against PRELID1, MCU, EMRE and compared to control (Scr). ATP5A is used as a loading control. **E**, Representative traces and quantification of [Ca^2+^]_mt_ transients upon histamine (Hist) stimulation in PRELID1-silenced HeLa mt-AEQ cells (mean ± SEM; n= 5); student’s t-test (***p <0.001). **F**, Domain structure of EFHD1 highlighting two evolutionarily conserved EF-hand domains (EF1 and EF2) and a coiled-coil domain (CCD). **G**, Immunoblot analyses of EFHD1 protein level in whole cell lysates from shRNA-mediated EFHD1 knockdown (sh1-5) and control (pLKO) cells. *CDS*, coding sequence. **H**, **I**, Representative traces and quantification of [Ca^2+^]_mt_ transients upon histamine (Hist) stimulation in HeLa mt-AEQ cells expressing either EFHD1-targeting sh-RNAs (H) or si-RNAs (I) (mean ± SEM; n> 24 for H and n> 26 for I); one-way ANOVA with Dunnett’s multiple comparisons test (*p< 0.05, ***p< 0.001, ****p< 0.0001). **J**, Average time courses and quantification of histamine-induced [Ca^2+^]_mt_ (left panel) and [Ca^2+^]_cyt_ responses (right panel) in si-EFHD1 HeLa cells. Peak and area under the curve (AUC) are calculated for the first 60 s of histamine (Hist) stimulation (mean ± SEM; Scr, n= 60; si1-EFHD1, n= 65; si2-EFHD1, n= 63); one-way ANOVA with Dunnett’s multiple comparisons test (*p< 0.05, **p< 0.01; *ns*, not significant). **K**, Representative traces and quantification of [Ca^2+^]_mt_ (left panel) and [Ca^2+^]_cyt_ (right panel) responses upon 1.5 mM Ca^2+^ induced SOCE activation in si-EFHD1 HeLa mt-AEQ cells (mean ± SEM; n≥ 10); one-way ANOVA with Dunnett’s multiple comparisons test (*p< 0.05, **p< 0.01; *ns*, not significant). **L**, Representative traces and quantification of [Ca^2+^]_mt_ transients upon histamine (Hist) stimulation in HeLa mt-AEQ cells transfected with a C-terminal V5-tagged WT or mutant EFHD1 in both EF-hand domains (EFHD1_EF1+2_; EF1: D231A, E242K; EF2: D421A, E432K). Indel shows immunoblot analysis of EFHD1 protein expression. ACTIN is used as a loading control (mean ± SEM; n> 24); one-way ANOVA with Dunnett’s multiple comparisons test (*p< 0.05). See also **Figures S2-S3.**

As a third candidate to test we chose EFHD1, a poorly characterized EF-hands mitochondrial protein (**Fig. 4F**), whose role in mt-Ca^2+^ homeostasis remains controversial (Meng *et al*, 2023; Hou *et al*, 2016; Eberhardt *et al*, 2022). EFHD1 was recently shown to bind MCU in clear cell renal cell carcinoma (ccRCC) (Meng *et al*, 2023) and it therefore provides a great example of the predictive power of our resource. Indeed, we identified EFHD1 as a high-confidence and common interactor of all three baits suggesting a strong functional connection to MCUC. To test this hypothesis, we first used aequorin as a luminescent Ca^2+^ sensor to quantify [Ca^2+^]_mt_ and [Ca^2+^]_cyt_ transients in HeLa cells upon either stable or transient KD of EFHD1 by shRNA and siRNA treatment, respectively. Out of five distinct shRNAs targeting different regions of EFHD1 mRNA, two (sh1 and sh5) strongly decreased EFHD1 protein level (**Fig. 4G**) and caused a significant up-regulation of [Ca^2+^]_mt_ in response to histamine stimulation (**Fig. 4H**). Likewise, we observed higher [Ca^2+^]_mt_ upon transfection of HeLa cells with two distinct EFHD1-targeting siRNAs (si1, si2) compared to control (Scr) (**Fig. S3A** and **Fig. 4I**). In both experimental set-ups, histamine-stimulated [Ca^2+^]_cyt_ responses remained unaffected by silencing of EFHD1 (**Fig. S3B, C**). We corroborated these results by quantitative and simultaneous single-cell analyses of [Ca^2+^]_mt_ and [Ca^2+^]_cyt_ using a mitochondrial matrix-targeted RCaMP and Fura-2 AM, respectively, as fluorescent Ca^2+^ probes. The measurements were performed in the absence of extracellular Ca^2+^ to study the ER-to-mitochondria Ca^2+^ transfer without the involvement of SOCE. Histamine stimulation in si-EFHD1 but not Scr cells triggered a significant increase in both peak and area under the curve (AUC) for the [Ca^2+^]_mt_ response, without affecting [Ca^2+^]_cyt_ transients (**Fig. 4J**). Similar results were observed in the presence of 1.8 mM Ca^2+^ in the extracellular medium, when both agonist-induced ER Ca^2+^ release and SOCE were activated (**Fig. S3D**). To evaluate whether the histamine-stimulated mt-Ca^2+^ phenotype was due to EFHD1-dependent changes in the propagation of Ca^2+^ signals from the ER to the mitochondria, we also quantified [Ca^2+^]_mt_ and [Ca^2+^]_cyt_ transients upon opening of SOCE channels at the plasma membrane. To this goal, we pre-treated cells with thapsigargin, an inhibitor of the ER-localized Ca^2+^ ATPase (SERCA) pump, to deplete ER stores and activate SOCE. The addition of 1.5 mM of Ca^2+^ to the extracellular medium of si-EFHD1 HeLa cells resulted in [Ca^2+^]_mt_ transients of higher peak and amplitude compared to Scr, without a concomitant difference in [Ca^2+^]_cyt_ dynamics (**Fig. 4K**), consistent with the results obtained with histamine stimulation. Next, to understand whether the regulation of mt-Ca^2+^ uptake by EFHD1 was dependent on Ca^2+^ binding, we generated HeLa cells expressing either wildtype (WT) EFHD1 or a mutant in both EF-hand domains (EFHD1_EF1+2_). Although the overexpression of WT EFHD1 did not affect histamine-stimulated [Ca^2+^]_mt_ response, EFHD1_EF1+2_ cells phenocopied the effect of EFHD1 KD, causing an increase in [Ca^2+^]_mt_ (**Fig. 4L**). This suggests that binding of Ca^2+^ to EFHD1 is required to exert an inhibitory function on MCUC.

Altogether, we corroborate a role for MICU3 in the modulation of MCUC activity in HEK293 cells, whereby all MICUs co-exist; we validate PRELID1 as a novel molecular link between membrane phospholipid metabolism and [Ca^2+^]_mt_ homeostasis; we identify EFHD1 as a negative modulator of mt-Ca^2+^ uptake.

### EFHD1 is an IMS protein that regulates MCUC activity through MICU1 and affects the viability of breast and cervical cancer cells

To understand the mechanism of EFHD1-dependent inhibition of mt-Ca^2+^ uptake, we measured Ca^2+^ transients in mitochondria of digitonin-permeabilized mt-AEQ HeLa cells upon EFHD1 KD. The addition of exogenous Ca^2+^ triggered a dramatic increase of [Ca^2+^]_mt_ compared with control pLKO cells, and it was dependent on MCU activity given it was fully abrogated by pre-treatment with the ruthenium red derivative Ru360 (**Fig. 5A**). Because the impairment of IMM polarization and OXPHOS can also affect mt-Ca^2+^ uptake, we quantified mitochondrial bioenergetics in sh-EFHD1 HeLa cells (**Fig. 5B,C**). The oxygen consumption rate (OCR) of sh-EFHD1 cells at baseline, after treatment with the RCCV inhibitor oligomycin and the uncoupler carbonyl cyanide m-chlorophenylhydrazone (CCCP) was comparable to control cells, demonstrating that ATP-coupled respiration was unaffected and membrane potential was preserved (**Fig. 5B**). Consistently, when directly assessing glycolytic function, sh-EFHD1 and pLKO cells were indistinguishable (**Fig. 5C**). Next, we tested whether the inhibitory effect of EFHD1 on MCU-dependent Ca^2+^ uptake was due to a potential regulation of MCUC abundance or assembly, as previously shown for other uniporter components (Garbincius & Elrod, 2022). However, we found that neither stable nor transient silencing of EFHD1 in HeLa cells affected the expression of known MCUC subunits (**Fig. S4A**). Similarly, BN-PAGE of mitochondria from sh-EFHD1 HeLa cells did not reveal a significant change in the macromolecular profile of the complex compared to control cells (**Fig. S4B**), indicating that both stability and assembly of MCUC are preserved in the absence of EFHD1.

**Figure 5.**
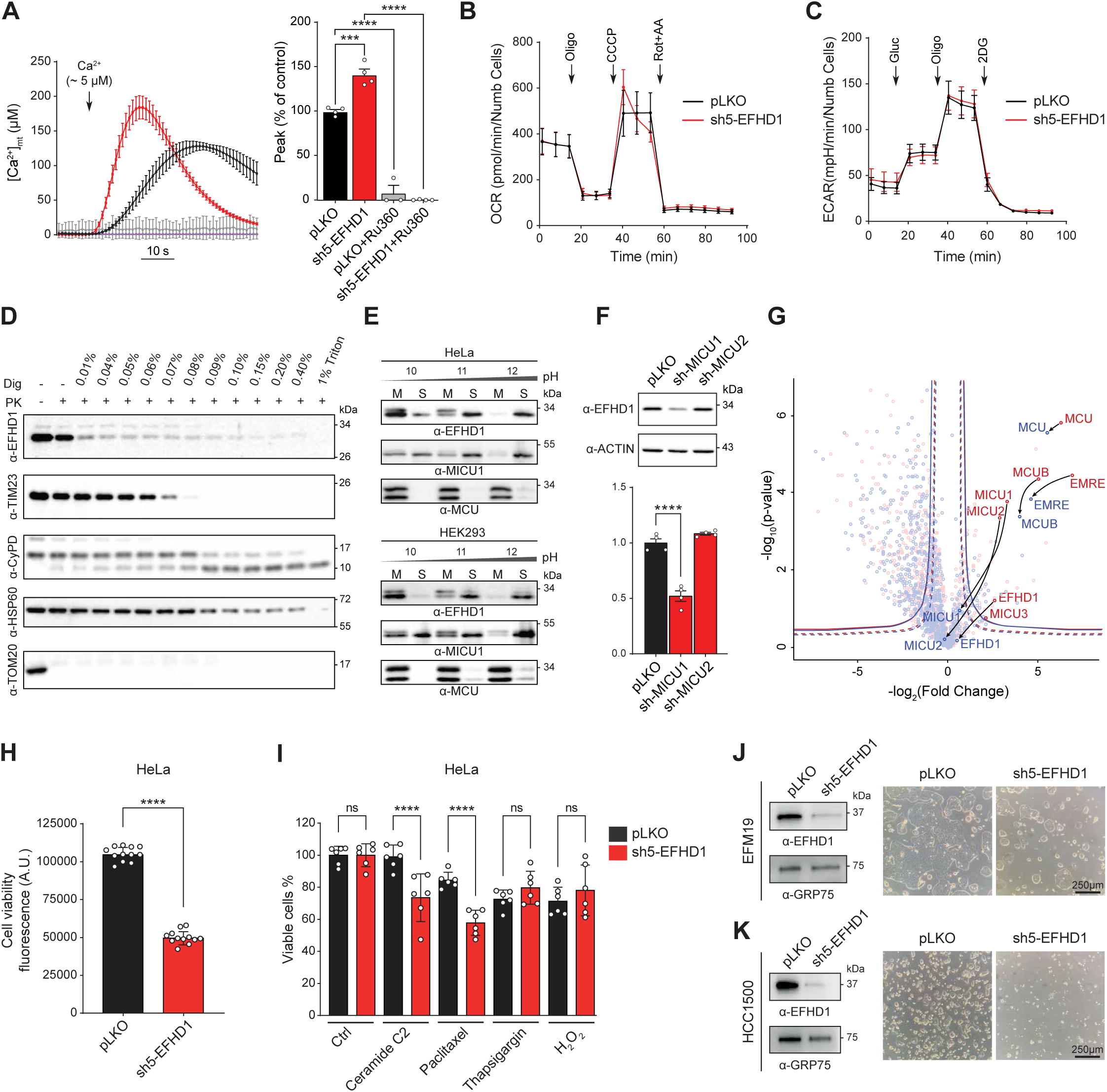
EFHD1 interaction with MCUC depends on MICU1 and affects viability of breast and cervical cancer cells. **A**, Quantification of [Ca^2+^]_mt_ upon addition of Ca^2+^ to permeabilized control (pLKO) and sh-EFHD1 mt-AEQ-HeLa cells (mean ± SEM; n= 4); one-way ANOVA with Dunnett’s multiple comparisons test (***p< 0.001, ****p< 0.0001). **B**, Normalized oxygen consumption rate (OCR) in control and sh-EFHD1 HeLa cells in response to oligomycin (Oligo, 1.5 µM), CCCP (1.5 µM), and Rotenone/Antimycin A (Rot/AA, 4 µM/2 µM), (mean ± SD; n≥ 10). **C**, Normalized extracellular acidification rate (ECAR) upon addition of glucose (Gluc, 10 mM), oligomycin (Oligo, 1.5 µM), and 2-deoxyglucose (2-DG, 100 mM), (mean ± SD; n≥ 10). **D**, Immunoblot analysis of EFHD1 in mitochondria from HeLa cells treated with Proteinase K (PK) at increasing digitonin (Dig) concentrations. TOM20, TIM23, Cyclophilin D (CyPD), and HSP60 are used as positive controls for OMM, IMS, and matrix proteins, respectively. **E**, Immunoblot analysis of EFHD1 in soluble (S) and membrane (M) fractions of mitochondria from HeLa and HEK293 cells after alkaline carbonate extraction at pH 10, pH 11, and pH 12. MICU1 and MCU are used as positive controls for membrane associated and integral membrane proteins, respectively. **F**, EFHD1 abundance in whole cell lysates from control, sh-MICU1, and sh-MICU2 HEK293 MCU-flag cells. ACTIN is used as a loading control (mean ± SEM; n= 4); one-way ANOVA with Dunnett’s multiple comparisons test (****p< 0.0001). **G**, Volcano plot of mitochondrial proteins enriched in MCU-TAP upon sh-MICU1 (blue) compared to pLKO control (red). Continuous and dashed lines indicate specific interaction partners defined based on a permutation-based FDR of either 0.05 or 0.10, respectively. **H**, Viability of HeLa cells after stable EFHD1 KD compared to control (pLKO) (mean ± SD; n= 12); student’s t-test (****p< 0.0001). **I**, Percentage of viable cells in HeLa pLKO and EFHD1 KD after treatment with apoptotic inducers (C2-ceramide, 40 µm; paclitaxel, 50 nM; thapsigargin, 500 nM; H_2_O_2_, 0.5 mM), compared to untreated cells (mean ± SD; n= 6); Fisher’s LSD test; ****p< 0.0001. **J**,**K**, Immunoblot analysis of EFHD1 (left) and representative images (right) of EFM19 (E) and HCC1500 (F) cells upon shRNA-mediated silencing of EFHD1. GRP75 is used as a loading control. See also **Figures S4-S5.**

To gain insights into the functional relationship between EFHD1 and MCUC, we characterized EFHD1 localization and topology. First, we performed a proteinase K (PK) assay on crude mitochondria isolated from HeLa cells (**Fig. 5D**). Proteins that are localized to the OMM and are exposed to the cytosol, such as TOM20, were immediately digested upon addition of PK, even in the absence of digitonin, while IMS-facing (TIM23) or matrix-localized (HSP60 and CypD) proteins were protected from PK until subsequent permeabilization of the OMM or IMM, respectively. EFHD1 showed a protein digestion profile comparable to that of TIM23 and consistent with an IMS localization. We obtained similar results when employing a PEGylation assay, in which OMM and IMM were selectively permeabilized by increasing concentrations of digitonin in the presence of maleimide functionalized polyethylene glycol (mPEG) (**Fig. S4C**). This reagent selectively adds a ∼5 kDa polyethylene glycol (PEG) polymer chain on free thiol groups of cysteines and is small enough to cross the OMM and directly react with IMS proteins. As it was reported for MICU1(Tsai *et al*, 2016), EFHD1 was pegylated even in the absence of digitonin, in contrast to the mitochondrial matrix protein, SOD2. Consistent with the lack of a predicted transmembrane domain, we found that EFHD1 was only associated with, but not spanning the IMM, because it was recovered in the soluble fractions of mitochondria from HeLa and HEK293 cells upon carbonate extraction, at both low and high pH, as MICU1 (**Fig. 5E**). The presence of EF-hand domains, EF-hand-dependent mt-Ca^2+^ uptake regulation, and an interaction with MCUC, together with evidence of an IMS localization, raised the hypothesis that EFHD1 functions is linked to other MICUs. Although, EFHD1 was unable to oligomerize through cysteine-mediated disulfide bonds (**Fig. S4D**) and to affect the formation of MICU1-MICU2 heterodimers, just like MICU2, it was cross-stabilized by MICU1 (**Fig. 5F**). Thus, we speculated that MICU1 could mediate the binding of EFHD1 to MCUC by affecting its intra-protein stability. Indeed, upon MICU1 silencing we found that the interaction between EFHD1 and MCU-tag was dramatically impaired, with a fourfold reduction in fold change and a loss of significance compared to control (**Fig. 5G**).

Since the sustained increase of [Ca^2+^]_mt_ has been shown to promote cell death, we speculated that high EFHD1 expression would protect cancer cells from pro-apoptotic triggers. As shown in **Fig. 5H**, the KD of EFHD1 in HeLa cells significantly decreased cell viability and sensitized cells to apoptotic inducers such as C2-ceramide and paclitaxel (**Fig. 5I**). We next set out to evaluate the relevance of EFHD1 more broadly in cancer biology. Indeed, expression profiling of a subset of tumor types, such as clear cell renal cell carcinoma (ccRCC) and colorectal cancer, have recently proposed the downregulation of EFHD1 as a prognostic factor and biomarker (Meng *et al*, 2023). However, a comprehensive assessment of its molecular signature in cancer cells and tissues is currently lacking. To this end, we analyzed publicly available cancer cell lines, primary tumors and patient datasets from the Cancer Cell Line Encyclopaedia (CCLE) of the Dependency Map Consortium (DepMap) (Ghandi *et al*, 2019), Gene Expression Omnibus of the National Centre for Biotechnology Information (NCBI-GEO), Cancer Genome Atlas (TCGA), Therapeutically Applicable Research to Generate Effective Treatments (TARGET), and Genotype-Tissue Expression (GTEx) repositories (Jiang *et al*, 2020). Compared to known components of MCUC such as MCU, EMRE, MICU1, MICU2 and MCUB, EFHD1 showed great variability in gene expression across 1450 cancer cell lines from 29 different lineages (**Fig. S5A**) but was consistently upregulated in breast, uterine, ovarian, and cervical cancer cells, both at the RNA and protein (**Fig. S5B**) level. Moreover, the expression of EFHD1 in primary breast tumor biopsies was significantly higher compared to the adjacent healthy tissues (**Fig. S5C**), and breast cancer patients with the highest EFHD1 expression exhibited a decreased response to chemotherapy (**Fig. S5D**) and a lower probability of survival (**Fig. S5E**). Consistently, downregulation of EFHD1 in two breast cancer cell types, HCC1500 and EFM19, with the highest EFHD1 protein expression compared to the median of all CCLE lines (5.99-fold and 5.91-fold, respectively), resulted in a dramatic decrease in cell viability and proliferation (**Fig. 5J,K**) that was not accompanied by a change in invasive potential (**Fig. S5F**).

### Genetic perturbation of MCUC greatly remodels the MCU protein network

Mapping of the MCUC network at physiological conditions clearly highlighted that the molecular composition of MCUC and the functional associations between mitochondria and Ca^2+^ are far more complex than previously anticipated (Sancak *et al*, 2013). Moreover, to what extent genetic perturbations of MCUC impact function, structure, and regulation of mitochondrial processes has not been systematically assessed so far. To this goal, we sought to map the remodeling of MCUC interactome upon LOF of EMRE, MICU1, MICU2, and MCUB. We generated isogenic MCU-tagged HEK293 cell lines stably expressing either short hairpin RNAs (shRNAs) targeting MICU1, MICU2, EMRE and MCUB, or pLKO as a control. As shown in **Fig. 6A**, we achieved an almost complete KD of each target. As previously reported (Kamer & Mootha, 2014), we observed that MICU1 KD resulted in the concomitant reduction of MICU2 and EMRE protein levels in whole cell lysates, whereas MICU1 expression was not affected by silencing of EMRE, MICU2 or MCUB. Instead, neither MICU2 nor MCUB downregulation altered the overall stability of other MCUC members. We then performed TAP-MS analyses upon tetracycline-inducible expression of StrepII-HA-His_6_ tagged MCU in stable pLKO and shRNA-expressing HEK293 cells in at least 4 biological replicates (**Table S4**). Principal component analysis showed great reproducibility across biological replicates and a clear separation between the interactome of MCU in control (pLKO) versus EMRE and MCUB KD, while clustering together with MICU1 and MICU2 KD, suggesting there are minimal changes to the MCU interaction network upon MICUs KD (**Fig. 6B**). By quantitative bait-prey co-enrichment analysis and local correlation of protein intensity profiles, we mapped PPIs between MCU and 245 mitochondrial proteins (**Table S4**). As expected, we observed that the interaction of MICU2 with MCU was dependent on MICU1 (Plovanich *et al*, 2013; Kamer & Mootha, 2014; Patron *et al*, 2014) and that silencing of EMRE resulted in the loss of MICUs (Sancak *et al*, 2013) as significant binding partners (**Fig. 6C**). Interestingly, although the MICU1-MCU interaction was not affected by MICU2 KD, MICU3 was not significantly co-enriched, suggesting that in absence of MICU2, the MICUs dimers are mostly formed by MICU1. On the contrary, the interaction of MCU with MCUB was always preserved, except upon silencing of MCUB, indicating that MCU and MCUB form a stable complex. Strikingly, the global analysis of all mapped MCUC interactomes spotlighted a dramatic remodeling of PPIs upon LOF of EMRE and MCUB (**Fig. 6D, E**). Downregulation of EMRE resulted in more than 10-fold lower prey recovery. However, although loss of EMRE leads to non-functional MCU channels (Sancak *et al*, 2013; Liu *et al*, 2020; König *et al*, 2016), we also identified novel and significantly enriched interactions between MCU and proteins of the TCA cycle that are known to be regulated by Ca^2+^, for example the pyruvate dehydrogenase complex (**Fig. 6F**). It is tempting to speculate that such PPIs could mediate a compensatory metabolic response aimed at preserving mitochondrial energy production. This could explain why, despite both MCU and EMRE knockout (KO) mouse models displaying a complete loss of mt-Ca^2+^ uptake, only MCU KO mice showed impaired exercise performance (Liu *et al*, 2020; Pan *et al*, 2013). In the opposite direction, MCUB KD, which leads to increased [Ca^2+^]_mt_, neither affected MCUC protein stability nor the interaction of EMRE and the MICUs with MCU. However, it substantially remodeled the MCU interactome, mostly by allowing the gain of novel interactions. These involved mitochondrial proteins that regulate stress response pathways, cell death, and mitochondrial translation. To corroborate these results, we tested whether a wider protein interaction network would also be reflected in the formation of MCU-containing complexes with higher MW upon sh-MCUB. Blue native-polyacrylamide gel electrophoresis (BN-PAGE) analysis of mitochondria isolated from pLKO MCU-tagged HEK293 cells identified the expected bands around 480 kDa and above 720 kDa and showed that silencing of MICU1 or EMRE caused a shift toward a lower MW (Plovanich *et al*, 2013; Sancak *et al*, 2013), while MICU2 LOF did not affect the overall assembly (Plovanich *et al*, 2013) (**Fig. 6G**). Instead, stable (shRNA) and transient (siRNA) MCUB KD resulted in the shift of MCU-containing complexes toward a higher MW, both in HEK293 (**Fig. 6G**) and HeLa cells (**Fig. 6H**). We observed the same phenotype in mitochondria isolated from the brain of MCUB KO mice, where we and others (Samaras *et al*, 2020; Hansen *et al*, 2024) could detect a clear expression of MCUB (**Fig. S6A-C**). Moreover, two-dimensional blue native/SDS analysis of whole brain mitochondria from MCUB KO confirmed that MCU-containing complexes were shifted toward a higher MW (**Fig. 6I**). In summary, our results suggest that the extent to which MCU engages in PPIs within the organelle is determined by channel activity and [Ca^2+^]_mt_ and that MCUB can act as a protein barrier **(Fig. 6J)**, by preventing PPIs between MCU and several mitochondrial complexes and pathways.

**Figure 6.**
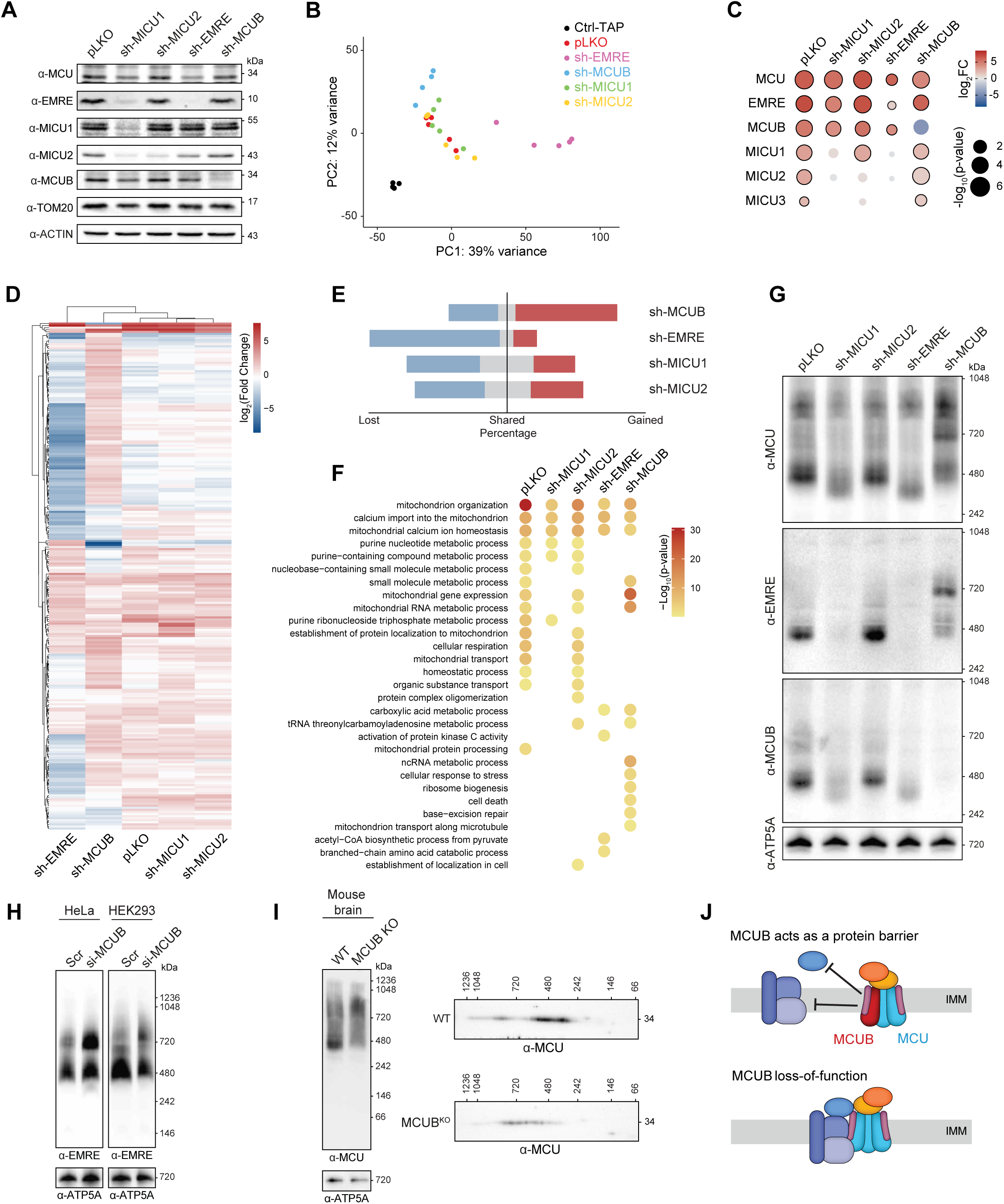
Remodeling of the MCU interactome upon genetic perturbation of the uniporter complex. **A**, Immunoblot analysis of known MCUC components in Flp-In T-REx HEK293 cells expressing MCU as a bait and infected with either control (pLKO) or shRNAs targeting MICU1, MICU2, EMRE and MCUB. ACTIN and TOM20 are used as loading controls. **B**, Principal Component (PC) Analysis of TAPs from MCU-tagged and parental cell lines. **C**, Quantitative bait-prey co-enrichment analysis of the previously known MCUC components upon genetic perturbation. Dots with a black border indicate significant interactions based on FDR cutoff = 0.10. **D**, Heatmap showing differences in the Log_2_ fold change among MCU interactors across all tested conditions. Hierarchical clustering was performed using the Euclidean distance with complete linkage. **E**, Percentage of gained (red), shared (grey), and lost (blue) interactors in each condition compared to MCU-TAPs and calculated based on the union of TAPs from MCU pLKO and shRNAs. **F**, Pathway enrichment analysis of the significant interactors for pLKO and shRNAs conditions. **G**, BN-PAGE analysis of mitochondria isolated from Flp-In T-REx HEK293 cells expressing MCU-tagged and either pLKO or shRNAs. ATP5A is used as a loading control. **H**, BN-PAGE analysis of MCUC assembly in mitochondria isolated from control (Scr) and si-MCUB HeLa or HEK293 cells. ATP5A is used as a loading control. **I**, Blue Native/SDS PAGE analysis of MCUC in isolated mitochondria from WT and MCUB KO mouse brains. ATP5A is used as a loading control. **J**, Model of MCUB as a protein barrier: Incorporation of MCUB into the MCUC obstructs protein-protein interactions between MCU and various mitochondrial complexes and pathways. Loss-of-function of MCUB enhances the number of PPIs involving MCU and facilitates interaction with different mitochondrial functions. See also **Figure S6**, **Table S4** and **Table S5.**

## DISCUSSION

Proteins typically exert their function by functionally and physically interacting. To this end, several biochemical approaches have so far been exploited to map the protein interactome landscape of the uniporter in HEK293 cells (Antonicka *et al*, 2020; Sancak *et al*, 2013; Austin *et al*, 2022), mostly using MCU as a bait and upon its overexpression, a condition known to affect mitochondrial physiology and cell survival (De Stefani *et al*, 2011; D’Angelo *et al*, 2023; Mammucari *et al*, 2015). As a result, MCU, MCUB, EMRE, MICU1 and MICU2 were proposed to represent all the subunits of the so-called “holocomplex” in this cell type (Sancak *et al*, 2013). Compared to those analyses, we were able to expand the “holocomplex” by mapping 139 statistically significant PPIs between known members of the uniporter and an additional 89 mitochondrial proteins localized in all four sub-mitochondrial compartments. Key to our approach was the use of all three membrane-spanning subunits of the complex as baits and the integration of complementary computational strategies to systematically analyze MCUC copurifying proteins. Of utmost importance, our study provides the first snapshot of the MCUC signaling network at near-endogenous conditions. Besides being well supported by recent literature and providing novel insights, our results identify proteins that regulate the MCUC activity as well as functional associations between MCU-mediated mt-Ca^2+^ uptake and numerous mitochondrial complexes and pathways. As an example, we recovered functional interactions between MCU, EMRE, the YME1L proteolytic hub, and the m-AAA protease AFG3L2 that are required for MCUC biogenesis and assembly (König *et al*, 2016; Tsai *et al*, 2022, 2017). We also mapped several molecular links between the MCUC, energy production, and cell death activation, mitochondrial functions that are known to be regulated by calcium (Rizzuto *et al*, 2012). We identified as MCUC interactors proteins involved in mt-DNA maintenance and replication, mRNA transcription and protein translation, suggesting for the first time a role for mt-Ca^2+^ on the organelle’s transcriptional and translational control. Similarly, the MCUC network pointed to a tight link between [Ca^2+^]_mt_ homeostasis and lipid metabolism, as it included several enzymes of the beta-oxidation pathway (e.g. ECHS1, ACACB, ACOT7). Interestingly, the impairment of mt-Ca^2+^ uptake was already associated with the rewiring of energy production from glycolysis to fatty acid oxidation in the skeletal muscle (Kwong *et al*, 2018; Gherardi *et al*, 2018; Huo *et al*, 2023), but the molecular mechanisms underlying such metabolic flexibility remain unknown.

The high coverage and sensitivity of our approach were further corroborated by the identification of MICU3 as a *bona fide* component of MCUC in HEK293 cells. None of the previous PPIs analyses (Sancak *et al*, 2013; Antonicka *et al*, 2020; Austin *et al*, 2022) recovered MICU3 within the set of MCU copurifying proteins, possibly due to its low expression level in HEK293 cells (Morgenstern *et al*, 2021), rather than its tissue-specific expression (Plovanich *et al*, 2013; Patron *et al*, 2019). Our MCUC interactome represents also a great resource to discover additional regulatory mechanisms of mt-Ca^2+^ signaling. For example, we identified EFHD1 as a high-confidence binding partner of all three baits. So far, EFHD1 has been linked to mitochondrial flashes (Hou *et al*, 2016; Eberhardt *et al*, 2022), ROS signaling (Eberhardt *et al*, 2022), bioenergetics (Ulisse *et al*, 2020), pro-B immune cell development and maturation (Stein *et al*, 2017), as well as cell survival and proliferation (Meng *et al*, 2023). However, its function in mitochondria and the mechanisms responsible for such a pleiotropic effect in different cell and tissue types remain poorly understood. Its involvement in [Ca^2+^]_mt_ homeostasis has been explored in previous reports but with some inconsistencies (Hou *et al*, 2016; Meng *et al*, 2023; Eberhardt *et al*, 2022). Whereas EFHD1 silencing in HeLa cells was not associated to a defect in [Ca^2+^]_mt_ upon either osmotic stress or histamine stimulation (Hou *et al*, 2016), a slight decrease in [Ca^2+^]_mt_ was observed after EFHD1 KO in cardiomyocytes (Eberhardt *et al*, 2022), potentially due to a drop in the mitochondrial membrane potential. On the other hand, the overexpression of EFHD1 in ccRCC was found to significantly reduce [Ca^2+^]_mt_ (Meng *et al*, 2023). To solve this conundrum, we performed complementary and quantitative measurements of intracellular Ca^2+^ dynamics upon either stable or transient KD of EFHD1 in HeLa cells, employing both luminescence– and fluorescence-based Ca^2+^ assays. Collectively, our results indicate that EFHD1 functions as an inhibitor of MCU-mediated [Ca^2+^]_mt_ uptake. This role is neither due to changes in upstream Ca^2+^ signaling pathways nor in oxidative metabolism, but seems to be dependent on its interaction with the MCUC, observation supported by its cross-stabilization with MICU1. This possibly allows EFHD1 to sense changes in [Ca^2+^]_mt_ and regulate MCUC activity accordingly, as performed by other EF-hand-containing proteins such as the MICUs (Csordás *et al*, 2013; Plovanich *et al*, 2013; Patron *et al*, 2019). Importantly, the heterogeneous expression pattern of EFHD1 among human tissues and cancer cell lines could also explain why EFHD1 loss– or gain-of-function would affect [Ca^2+^]_mt_ homeostasis to different extents in different cell types. Accordingly, in HeLa cells that already express EFHD1 at a high level, we failed to observe an alteration in [Ca^2+^]_mt_ upon EFHD1 overexpression. Instead, its overexpression in cells like ccRCC that show 30-fold lower level of endogenous EFHD1 compared to HeLa cells was found to significantly reduce [Ca^2+^]_mt_ (Meng *et al*, 2023). As a selective determinant of cell survival and cancer progression, EFHD1 surely represents an important therapeutic target for further investigations.

One of the most characteristic features of MCUC is the deep plasticity to adapt to and compensate mitochondrial dysfunctions. Indeed, the cross-regulation of protein level and stability among its components represents one of the main mechanisms behind compensatory adaptation observed upon chronic loss or gain of mt-Ca^2+^ uptake in cells and mouse models (Pan *et al*, 2013; Liu *et al*, 2020). For example, the reduction of [Ca^2+^]_mt_, by increasing MCUB:MCU (Huo *et al*, 2020; Lambert *et al*, 2019) or MICU1:MCU ratio (Paillard *et al*, 2022), allows to protect mitochondria from Ca^2+^ overload and associated cell death. Conversely, enhancing uniporter activity counteracts the bioenergetic deficit due to OXPHOS impairment and mitochondrial cardiomyopathies upon TFAM deletion (Balderas *et al*, 2022), as well as sustained adaptive thermogenesis in brown adipose tissue upon adrenergic stimulation (Xue *et al*, 2022). To gain insights into the remodeling of MCUC interactome upon genetic perturbation, we also performed TAPs followed by LC-MS/MS analysis from MCU-tagged cells where we stably silenced MICU1, MICU2, EMRE or MCUB. Our integrative analysis of the MCUC PPIs upon loss– or gain-of-function in mt-Ca^2+^ uptake, demonstrates that the MCU interactome is a flexible and adaptable network to perturbations and spotlights MCUB as a protein barrier. MCUB was initially described as a dominant negative regulator of [Ca^2+^]_mt_, due to critical substitution in its highly conserved transmembrane domain, which could impact the permeation of Ca^2+^ through the channel (Raffaello *et al*, 2013). We observed that silencing of MCUB led to a more extended MCUC network and was accompanied by an increased formation of high MW complexes of MCU, both *in vitro* and *in vivo*. These findings are also consistent with recent evidence showing that MCUB expression in cardiomyocytes leads to a reduction in MCUC size and disrupts its interaction with MICU1 and MICU2 (Lambert *et al*, 2019; Huo *et al*, 2020). However, our results indicate that the formation of MCU complexes with higher MW is not simply due to changes in PPIs between the known MCUC members, but to a wider remodeling of the MCU interactome.

In summary, our MCUC interaction map under physiological conditions and genetic perturbation of the different MCUC components, represents a great tool to (1) identify novel candidate proteins and pathways involved in [Ca^2+^]_mt_ signal transduction cascades; (2) provide new insights into both players and mechanisms regulating tissue-specific MCUC activity; (3) uncover novel genetic underpinnings and pharmacological targets to develop new therapeutical strategies for several human diseases.

## MATERIALS AND METHODS

### Cell lines

HeLa and HEK293 cells were cultured in Dulbecco’s modified Eagle’s medium (DMEM) supplemented with 10% FBS. HeLa cells stably expressing a mitochondrial matrix-targeted GFP-aequorin (mt-AEQ) (Arduino *et al*, 2017) were cultured in DMEM supplemented with 10% FBS and 100 µg/mL geneticin (Thermo Fisher Scientific, 10131027). EFM19 cells (German Collection of Microorganisms and Cell Cultures GmbH) and HCC1500 cells (American Type Culture Collection) were cultured in RPMI1640 media supplemented with 10% FBS. All cell lines were kept at 37°C in an incubator with 5% CO_2_.

The Flp-In T-REx system was used to generate isogenic, stable HEK293 cell lines exhibiting tetracycline-inducible expression of tagged bait proteins for tandem affinity purification (TAP). The open reading frame (ORF) of each bait protein (MCU, MCUB and EMRE) was cloned into the pcDNA5/FRT/TO Flp-In expression vector, in frame with a C-terminal StrepII_2_-HA-His_6_-tag and under the control of a tetracycline-regulated CMV/TetO2 promoter. The resulting MCU, EMRE, or MCUB Flp-In expression vectors were co-transfected with the Flp recombinase vector, pOG44, into Flp-In T-REx HEK293 cells, which contained a single integrated FRT site and stably expressed the lacZ-ZeocinTM fusion gene (pFRT/lacZeo) and the tetracycline repressor (pcDNA6/TR). This allowed the targeted integration of each bait protein at a single transcriptionally active genomic locus that is the same in every cell line. Flp-In T-REx HEK293 cells were grown in DMEM with 10% FBS, containing blasticidin (15 µg/mL) and Zeocin (100 µg/mL). Stable transfectants were selected and maintained in DMEM with 10% FBS containing blasticidin (15 µg/mL) and 100 µg/mL hygromycin. The expression of tagged bait proteins was induced by the addition of tetracycline (1 µg/ml) to a growth medium lacking hygromycin and blasticidin, 24 h prior to harvest.

Human cell lines stably expressing sh-RNAs for gene-specific knockdown (KD) were generated as previously described (Perocchi *et al*, 2010). The following sh-RNA constructs were used for MCU (sh-MCU, TRCN0000133861); EMRE (sh-EMRE, TRCN0000145067); MICU1 (sh-MICU1, TRCN0000053370); MICU2 (sh-MICU2 TRCN0000055848); MICU3 (sh1-MICU3, TRCN0000056083; sh2-MICU3, TRCN0000056084; sh3-MICU3, TRCN0000056085; sh4-MICU3, TRCN0000056086; sh5-MICU3, TRCN0000056087); EFHD1 (sh1-EFHD1, TRCN0000056183; sh2-EFHD1, TRCN0000056184; sh3-EFHD1, TRCN0000056185; sh4-EFHD1, TRCN0000056186; sh5-EFHD1, TRCN0000056187); and MCUB (sh-MCUB, TRCN0000128550).

Infected HeLa cells were selected with 2 µg/mL puromycin. EFM19 and HCC1500 cells were infected with pLKO and sh5-EFHD1 viruses and selected with 1 µg/mL puromycin. The same number of cells for each condition (50,000 for EFM19; 25,000 for HCC1500) were seeded in a 3 cm culture plate and imaged 10 days after seeding, with media changes every 48 h. For siRNA-mediated KD, 50 nM of negative control scramble or targeting siRNAs (Scr: MISSION siRNA #1 SIC001; si1-EFHD1: SASI_Hs01_00228164; si2-EFHD1: SASI_Hs01_00228165, si-MCUB: 5’-AUACUACCAGUCACACCAU-3’, si-PRELID1: 5’-CCCGAAUCCCUAUAGCAAA-3’) were transfected using Lipofectamine RNAiMAX (Thermo Fisher Scientific, 13778075). Assays were normally carried out 48 h after transfection, unless noted otherwise. For the exogenous expression of MICU3 and EFHD1 ORFs in human cells, MICU3 (HsCD00296366) and EFHD1 (HsCD00719312) WT clones without a STOP codon in pDONR221 were purchased from DNASU, whereas MICU3 and EFHD1 EF-hand mutants were synthesized in the pUC57 vector (Thermo Fisher Scientific, SD0171) and cloned into a pDONR221 vector with the following primers: attB1-FW-MICU3, GGGGACAAGTTTGTACAAAAAAGCAGGCTTAGCCACCATGGTGGCTCGAGGGCT; attB2-RV-MICU3, GGGGACCACTTTGTACAAGAAAGCTGGGTTTCTGCTGTGAAGTTCTTTCTTCAGG; attB1-FW-EFHD1, GGGGACAAGTTTGTACAAAAAAGCAGGCTTAGCCACCATGGCCAGTGAGGAGCTG; attB2-RV-EFHD1, GGGGACCACTTTGTACAAGAAAGCTGGGTTTGTATTGAAGTTGGCCTTGAGTTT. Constructs were then cloned either into pcDNA-DEST40 Vector (Thermo Fisher Scientific, 12274015) or pLX304 for expression in mammalian cells in frame with a C-terminal V5-His_6_ and V5 tag, respectively, by gateway cloning according to manufacturer instructions. Lentivirus production and infection were performed according to guidelines from the Broad RNAi Consortium and infected cell lines were selected 48 h post-transduction with the respective selection markers. Transient protein expression was performed using X-tremeGENE^TM^ HP DNA transfection reagent following the manufacturer’s instructions for a 1:3 DNA/reagent ratio. Assays were normally carried out 48 h after transfection, unless noted otherwise.

### Immunoblot analysis

To monitor endogenous and overexpressed proteins, cells or isolated mitochondria were lysated in RIPA-buffer (150 mM NaCl, 50 mM Tris, 1 mM EGTA, 1% NP40, 0.1% SDS); after 30 min of incubation on ice, the lysates were centrifuged at 15,000x*g* for 10 min to remove debris. 20 µg of total proteins were loaded, according to BCA quantification (Thermo Fisher Scientific, 23225) for each lane. Proteins were reduced with Laemmli buffer (Bio-Rad, 1610747) supplemented with 2.75 mM β-mercaptoethanol (Thermo Fisher Scientific, 21985-023) and denatured for 5 min at 90° C, unless otherwise specified. To visualize MICU1-MICU2 or MICU1-MICU3 heterodimers, protein samples were denatured at 70°C for 10 min in LDS Sample Buffer (Thermo Fisher Scientific, NP0007) with or without 100 mM Dithiothreitol (+DTT and –DTT, respectively), as previously performed (Patron *et al*, 2014). Proteins were separated by SDS-PAGE electrophoresis in 12% or 14% acrylamide gels, and transferred on nitrocellulose membranes (Amersham, 10600021) by semi-dry electrophoretic transfer (Bio-Rad). Accordingly with primary antibody datasheets, blots were blocked 1 h at room temperature (RT) with 5% non-fat dry milk (Carl Roth, T145.2) or 5% bovine serum albumin (BSA, Sigma-Aldrich, A7906) in TBS-Tween (0.5 M Tris, 1.5 M NaCl, 0.01% Tween) solution and incubated over night at 4°C with primary antibodies. Horseradish peroxidase-conjugated secondary antibodies (Bio-Rad, 1706515 or 1706516), diluted in TBS-Tween containing 0.5% BSA, were incubated for 1 h at RT followed by detection by chemiluminescence (Amersham, RPN2236). The expression level of specific proteins was detected by immunoblot analysis with the following antibodies: HA (BioLegend, MMS-101R), V5 (Thermo Fisher Scientific R96025), ACTIN (Sigma-Aldrich, A2228), MAIP1 (König *et al*, 2016), ATP5A (Abcam MS507), Lamin (Santa Cruz, sc-6217), VDAC (Abcam, ab14734), EFHD1 (Sigma-Aldrich HPA056959), HSP60 (R&D System MAB1800), SOD2 (Antibody Verify Inc. AAS29585C), TIM23 (BD bioscience, 611222), CyPD (Abcam, ab110324), GRP75 (Santa Cruz, sc-133137), MCU (Sigma-Aldrich, HPA016480), EMRE (Santa Cruz, sc-86337), MCUB (Santa Cruz, sc-163985), MICU1 (Sigma-Aldrich, HPA037479), MICU2 (Sigma-Aldrich, HPA045511), MICU3 (Sigma-Aldrich, HPA024048), TOM20 (Abcam, ab56783). Densitometry analysis of protein bands was performed with ImageJ by subtracting background signal and normalizing the area of each peak intensity to actin.

### Tandem affinity purification

Isogenic, parental Flp-In T-REx HEK293 cells stably expressing tagged MCU, MCUB, or EMRE as baits, as well as Flp-In T-REx HEK293 cells expressing tagged MCU with stable knockdown of either MICU1, MICU2, EMRE, or MCUB were expanded into two expression plates (245×245 mm, NUNC, 9407400) to obtain roughly 250 million cells. For TAP, each cell pellet was resuspended in 2 mL lysis buffer (50 mM HEPES-KOH pH 7.4, 150 mM KCL, 5 mM EGTA, 5% Glycerol, 3% Digitonin) and incubated at 4°C in rotation for 30 min. Lysates were centrifuged at 10000x*g* for 5 min at 4°C and the supernatants were incubated for 20 min at RT on a rolling shaker with 50 µL of a 500 mM HEPES solution at pH 8.0 containing avidin (10 µM final concentration). In the meantime, 150 µL of streptavidin resin was washed three times with 500 µL washing buffer (50 mM HEPES-KOH pH 7.4, 150 mM KCl, 5 mM EGTA, 5% Glycerol, 0.02% Digitonin), combined with the supernatant, and incubated 45 min at 4°C in rotation. Resins were then washed three times, before eluting bound proteins with 250 µL of elution buffer (50 mM HEPES-KOH pH 8, 150 mM KCl, 5 mM EGTA, 5% Glycerol, 10 mM Biotin), incubating on a rolling shaker at 900 rpm for 5 min at RT. The elution step was repeated 4 times and a total of 1 mL eluate was collected. Next, 50 µL of Ni-NTA resin was washed three times with 500 µL of washing buffer and incubated with the eluate for 45 min at 4°C in rotation. The resin was then collected by quick spin and washed three times with 500 µL of washing buffer followed by two washing steps in washing buffer without detergent. Purified complexes were eluted through three steps of incubation with 35 µL of 1% RapiGest on a rolling shaker for 10 min at 900 rpm at RT. Eluates were pulled and subjected to acetone precipitation overnight. The next day, samples were centrifuged at 20000x*g* for 30 min at 4°C, the supernatant was removed, and the pellet was stored at –80°C for LC-MS/MS analysis.

### Sample preparation for LC-MS/MS analysis

Eluates were resuspended in 50 µL denaturation buffer (6 M urea, 2 M thiourea, 10 mM HEPES pH 8.0, 10 mM DTT) for 30 min at RT before adding alkylation agent (55 mM iodoacetamide) and incubating at RT in the dark for 20 min. Proteins were digested at RT for 3 h by adding LysC at 1:100 ratio of enzyme:protein, before diluting the sample with 50 mM ammonium bicarbonate to reach an urea concentration of 2 M. Subsequently, samples were digested at RT overnight by adding Trypsin at 1:100 ratio of enzyme:protein. The next day, the digestion was stopped by acidifying the sample with 10 µL of 10% trifluoroacetic acid and the final peptides were cleaned up using SDB-RPS StageTips as described (Kulak *et al*, 2014). MS analysis was performed using Q Exactive HF mass spectrometers (Thermo Fisher Scientific, Bremen, Germany) coupled on-line to a nanoflow ultra-high-performance liquid chromatography instrument (Easy1000 nLC, Thermo Fisher Scientific). Peptides were separated on a 50 cm long (75 μm inner diameter) column packed in-house with ReproSil-Pur C18-AQ 1.9 μm resin (Dr. Maisch GmbH, Ammerbuch, Germany). Column temperature was kept at 50°C. Peptides were loaded with buffer A (0.1% (v/v) formic acid) and eluted with a nonlinear gradient of 5-60% buffer B (0.1% (v/v) formic acid, 80% (v/v) acetonitrile) at a flow rate of 250 nL/min. Peptide separation was achieved by 120 min gradients. The survey scans (300-1650 m/z, target value = 3E6, maximum ion injection times = 20 ms) were acquired at a resolution of 60000 followed by higher-energy collisional dissociation (HCD) based fragmentation (normalized collision energy = 27) of up to 15 dynamically chosen most abundant precursor ions. The MS/MS scans were acquired at a resolution of 15000 (target value = 1E5, maximum ion injection times = 60 ms). Repeated sequencing of peptides was minimized by excluding the selected peptide candidates for 20 s.

### MS data processing and visualization

Data analysis was carried out using the MaxQuant software package 1.5.5.2. The false discovery rate (FDR) cutoff was set to 1% for protein and peptide spectrum matches. Peptides were required to have a minimum length of 7 amino acids and a maximum mass of 4600 Da. MaxQuant was used to score fragmentation scans for identification based on a search with an initial allowed mass deviation of the precursor ion of a maximum of 4.5 ppm after time-dependent mass calibration. The allowed fragment mass deviation was 20 ppm. Fragmentation spectra were identified using the UniprotKB *Homo sapiens* database, combined with 245 common contaminants by the integrated Andromeda search engine. Enzyme specificity was set as C-terminus to arginine and lysine, also allowing cleavage before proline, and a maximum of two missed cleavages. Carbamidomethylation of cysteine was set as fixed modification and N-terminal protein acetylation as well as methionine oxidation as variable modifications. Both ‘label-free quantification (MaxLFQ)’ with a minimum ratio count of 1 and ‘match between runs’ with standard settings were enabled. Protein copy number estimates were calculated using the iBAQ algorithm, in which the sum of all tryptic peptides intensities for each protein is divided by the number of theoretically observable peptides. The mass spectrometric data have been deposited via PRIDE (Vizcaíno *et al*, 2013) to the ProteomeXchange Consortium under the accession number PXD040893.

Basic data handling, normalization, statistics, and annotation enrichment analysis was performed with the Perseus software package (1.6.15.0 release) (Tyanova *et al*, 2016), R (4.1.2 release) and GraphPad Prism (10.0 release). The label-free quantification (LFQ) module of the MaxQuant software (Cox & Mann, 2008; Cox *et al*, 2014) was used to define specific proteins that were quantitatively enriched with a given bait over all measured samples. Protein groups were filtered removing hits to the reverse decoy database and proteins only identified by modified peptides. Mitochondrial proteins were filtered using a curated list of 1276 proteins that integrates information from MitoCarta3.0 (Rath *et al*, 2021), Uniprot annotated mitochondria proteins, and IMPI (Smith & Robinson, 2019). It was required that each protein was quantified in at least two biological replicates from TAPs of each cell line to be considered for analysis. Protein LFQ intensities were log-transformed and missing values were imputed by values sampled from a normal distribution shifted 1.8 standard deviation and with a width 0.3 standard deviations from the distribution of all protein intensities within each sample as the background. Protein interactors that were quantitatively enriched with a given bait over Ctrl-TAP were selected by two-tailed Welch’s t test using the multiple volcano (Hawaii) plot option of Perseus (version 1.6.2.0) and a permutation-based FDR cutoff of 0.05 (Class A) and 0.10 (Class B) and S0=1 (Hein *et al*, 2015). Protein intensity profiles across each pair of bait and control TAPs were used to calculate a Pearson’s correlation coefficient between baits and preys (local correlation). A protein was defined as a specific interactor when having a correlation coefficient higher or equal than 0.6 based on the mean correlation of Class A interactors identified by volcano analysis (0.53) (Class C). Similarly, specific interactors were identified based on a Pearson’s correlation analysis across control, MCU-, EMRE– and MCUB-TAPs (global correlation) using a permutation-based FDR of 0.05 (Class D). For the remodelling of MCUC interaction network upon genetic perturbation, Class A, B, and C parameters were used to define MCU interactors. For sh-MCUB a permutation-based FDR cutoff of 0.025 and 0.05 for class A and B, respectively, and a correlation coefficient higher or equal to 0.8 for class C, were used.

### Protein-protein interaction network

MCU, MCUB, and EMRE protein-protein interactions (PPIs) were visualized using Cytoscape 3.10.0 (Shannon *et al*, 2003). For each MCUC interactor, biological function, number of transmembrane domains, sub-mitochondrial localization, and gene-disease associations were retrieved based on MitoCop (Morgenstern *et al*, 2021), literature-based manual curation, DeepTMHMM (Hallgren *et al*, 2022), MitoCarta 3.0 (Rath *et al*, 2021), and DisGeNET (Piñero *et al*, 2021). By uncharacterized proteins, we refer to proteins that lack single gene-based experimental evidence but whose function is inferred simply through large-scale analysis or sequence similarity. Protein expression level of MCUC interactors across 201 samples from 32 different tissue types of normal human individuals was obtained from GTEx (Jiang *et al*, 2020). The tissue numbers were reduced by removing some samples (Artery – Coronary, Esophagus – Mucosa, Minor Salivary Gland, Nerve – Tibial, Pituitary, and Skin) and by averaging tissues belonging to the same organ (arteries, colon segments, esophagus segments, and heart compartments), obtaining 21 tissues in total. Orthologs of MCUC interactors were identified across 120 eukaryotic species and one prokaryotic species (*E. coli*) used as outgroup and are common to CLIME (Li *et al*, 2014) and ProtPhylo (Cheng & Perocchi, 2015). Orthologs were defined using OBH (one-way best hit) as in{Formatting Citation}. The NCBI taxonomy database and the R package taxize were used to build the species tree. Hierarchical clustering was performed using the Euclidean distance with complete linkage. Significant interactors were analyzed using gProfiler’s GOSt tool (Raudvere *et al*, 2019). Significance was established using the algorithm gSCS as multiple testing correction method with a significance threshold of 0.05. Only GO biological process driver terms significant in at least one condition were taken in consideration.

### Measurements of Ca^2+^ transients

Measurement of extracellular Ca^2+^ clearance by mitochondria from digitonin-permeabilized HEK293 cells was performed using the membrane impermeable Ca^2+^ indicator Calcium-Green-5N (Life technologies, C3737). Cells were harvested at a density of 500000 cells/mL in growth medium supplemented with 20 mM HEPES (pH 7.2/NaOH). Cells were collected by centrifugation at 300x*g* for 3 min at RT, re-suspended in extracellular-like media (145 mM NaCl, 5 mM KCl, 1 mM MgCl_2_, 10 mM Glucose, 10 mM HEPES, pH 7.4) containing 200 nM thapsigargin and incubated for 10 min under constant agitation at RT. Cells were then re-suspended in intracellular-like buffer (140 mM KCl, 1mM KH_2_PO_4_/K_2_HPO_4_, 1 mM MgCl_2_, 20 mM HEPES, 100 µM EGTA, pH 7.2/KOH), supplemented with 1 mM Na^+^-pyruvate, 1 mM ATP/MgCl_2_ and 2 mM Na^+^-succinate at a density of 2.5 x 10^6^ cells/mL and the plasma membrane was permeabilized by incubation with 60 µM digitonin for 5 min at RT under constant agitation. 100 µL of the cell suspension was seeded into a black 96-well plate (Perkin Elmer) and Calcium Green-5N fluorescence (excitation 506 nm, emission 531 nm) was monitored every 2 s at RT using a CLARIOstar microplate reader (BMG Labtech) after injection of 40 µM CaCl_2_. The MCU inhibitor Ru360 (10 µM) was used as a positive control.

Measurements of Ca^2+^ transients in HeLa cells were performed using the luminescence Ca^2+^ indicator aequorin as previously described (Arduino *et al*, 2017). Briefly, HeLa mt-AEQ cells infected with lentivirus carrying the specific shRNA were seeded in white 96-well plates (PerkinElmer, 6005181) at 25000 cells/well. After 24 h, aequorin was reconstituted with 2 µM native coelenterazine for 1-2 h at 37°C. For measurements of Ca^2+^ kinetics upon siRNA-mediated EFHD1 knockdown, cells were transfected using a transfection mix including Lipofectamine RNAiMAX transfection reagent (Thermo Fisher Scientific, 13778075), 0.5 µg of mt-AEQ (Rizzuto *et al*, 1992) or cytosolic aequorin (Brini *et al*, 1995) cDNAs, and either a final concentration of 50 nM siRNA. Mt-AEQ-based measurements of Ca^2+^-dependent light kinetics were performed upon 100 µM histamine stimulation. Light emission was measured either using the luminescence counter MicroBeta2 LumiJET Microplate Counter (PerkinElmer) or the PerkinElmer Envision plate reader at 469 nm every 0.1 and 1 s, respectively. For measurements of Ca^2+^ kinetics upon EFHD1 WT or EFHD1_(EF1+EF2)_ overexpression, cells were transfected using TransIT-2020 Transfection Reagent (Mirus Bio) with 1 ug of EFHD1 cDNA and 0.5 µg of mt-AEQ or cytosolic aequorin. Ca^2+^ kinetics upon store-operated Ca^2+^ entry (SOCE) were measured upon pre-treatment with the irreversible SERCA inhibitor thapsigargin (100 nM) for 10 min in Ca^2+^-free modified Krebs-Ringer Buffer (135 mM NaCl, 5 mM KCl, 1 mM MgCl_2_, 0.4 mM KH_2_PO_4_, 1 mM MgSO_4_, 20 mM HEPES, 600 μM EGTA, 10 mM glucose, pH 7.4 at 37°C) followed by perfusion with the same medium without thapsigargin but supplemented with 1.5 mM CaCl_2_. The measurement of mt-Ca^2+^ uptake in digitonin-permeabilized mt-AEQ HeLa cells was performed as previously described (Wettmarshausen *et al*, 2018). Briefly, HeLa cells stably expressing mt-AEQ were harvested at a density of 500,000 cells/mL in growth medium supplemented with 20 mM HEPES (pH 7.4/NaOH) and the aequorin was reconstituted by incubation with 2 µM native coelenterazine *n* (Biotium, BOT-10115-1) for 2.5 h at RT. Cells were then centrifuged at 300x*g* for 3 min and the pellet was re-suspended in an extracellular-like buffer (145 mM NaCl, 5 mM KCl, 1 mM MgCl_2_, 10 mM Glucose, 10 mM HEPES, pH 7.4) containing 200 nM thapsigargin and incubated for 20 min under constant agitation at RT. The cells were collected by centrifugation at 300x*g* for 3 min and the pellet was resuspended in an intracellular-like buffer supplemented with 1 mM Na^+^-pyruvate, 1 mM ATP/MgCl_2_ and 2 mM Na^+^-succinate. The cells were permeabilized for 5 min with 60 µM digitonin, collected by centrifugation for 3 min at 300x*g* and resuspended in intracellular-like buffer at a density of 900 cells/µL. Then, 90 µL of cell suspension was dispensed into a white 96-well plate (PerkinElmer). Cells were incubated for 5 min at RT and light signal was recorded at 469 nm every 0.1 s using a luminescence counter (MicroBeta2 LumiJET Microplate Counter, PerkinElmer) after injection of a 5 µM CaCl_2_ bolus. Ru360 (10 µM) was used as a positive control. All the luminescence signals were converted in [Ca^2+^] values accordingly to the Ca^2+^ response curve of aequorin, as previously performed (Brini *et al*, 1999).

For single cell measurement of Ca^2+^ transient, HeLa cells were treated with siRNA for 48 h before experiment. Cells were transiently transfected with mitochondrial matrix targeted RCaMP to measure changes in [Ca^2+^]_mt_ then plated on Poly-D-Lysin coated 25 mm coverslips (Thermo Fisher Scientific, 25CIR-1.5). To measure changes in the [Ca^2+^]_cyt_ cells were loaded with 2 μM Fura2AM (Moleculer probes, F-1221) in 2% BSA containing extracellular medium (ECM, 121 mM NaCl, 5 mM NaHCO_3_, 10 mM Na-HEPES, 4.7 mM KCl, 1.2 mM KH_2_PO_4_, 1.2 mM MgSO_4_, 2 mM CaCl_2_, and 10 mM glucose, pH 7.4) in the presence of 0.003% Pluronic F-127 (Thermo Fisher Scientific, P6867) and 150 μM sulfinpyrazone (Sigma Aldrich, S9509) for 15 min at 35°C. After dye-loading, cells were washed with fresh 0.25% BSA containing ECM with or without Ca^2+^ and transferred to the temperature-controlled stage (37°C) of an Olympus IX81 motorized inverted epifluorescence microscope fitted with a Hamamatsu ORCA-Flash 4.0v3 sCMOS camera, high speed excitation switching by Sutter Lambda 421 LED illuminator and UV-optimized Olympus UAPO/340 ×40/1.35NA oil immersion objective. For simultaneous measurements of [Ca^2+^]_cyt_ and [Ca^2+^]_mt_ Fura2 fluorescence was recorded at 340 and 380 nm, and RCaMP at 577 nm excitations, using dual-band Chroma 59022bs dichroic and 59022m emission filter. Image triplets were acquired every 0.33 s.

### Animals

C57BL/6n WT mice or C57BL/6n MCUB KO mice (Feno *et al*, 2021) were housed in a pathogen-free, temperature– and humidity-controlled animal facility on a 12:12 h light-dark cycle. Diet consisted of standard laboratory chow and double-distilled water. All animal procedures were in accordance with the European Community Council Directive for the Care and Use of Laboratory Animals (86/609/ECC) and German Law for Protection of Animals and were approved by the local authorities. All experiments were performed with female mice that were at least 3 months old.

### Isolation of functional mitochondria from cultured cells and mouse tissues

Mitochondria were isolated by nitrogen cavitation as previously described (Wettmarshausen & Perocchi, 2017). Briefly, HeLa and HEK293 cells were grown to confluency in a 600 cm^2^ cell culture plates. Culture medium was removed, and cells were rinsed with 30 mL PBS, scraped down and resuspended in 5 mL PBS. After 5 min of centrifugation at 600x*g* at 4°C, the cell pellet was resuspended in ice cold isolation buffer (IB; 220 mM mannitol, 70 mM sucrose, 5 mM HEPES-KOH pH 7.4, 1 mM EGTA-KOH, pH 7.4), supplemented with protease inhibitor (Sigma-Aldrich, 5056489001). Cell suspension was immediately subjected to nitrogen cavitation at 800 psi for 10 min at 4°C. Nuclei and intact cells were pelleted by centrifugation at 600x*g* for 10 min at 4°C. Supernatants were transferred into new tubes and centrifuged at 8000x*g* for 10 min at 4°C. The resulting pellet containing crude mitochondria was resuspended in 50-200 µL IB for further analyses. For mouse brain mitochondria isolation, the tissue was homogenized with two strokes at 300 rpm using a loose-fitting Teflon homogenizer followed by nitrogen cavitation at 800 psi for 10 min in ice cold isolation buffer supplemented with 0.5% fatty acid free BSA (Sigma-Aldrich, A7030) and protease inhibitor (Sigma-Aldrich, 5056489001). Nuclei and intact cells were pelleted by centrifugation at 600x*g* for 10 min at 4°C. Supernatants were transferred into new tubes and centrifuged at 12000x*g* for 10 min at 4°C. The centrifugation step was repeated, changing the buffer with IB without BSA. The final pellet was re-suspended in IB without BSA and stored on ice for further use. During the isolation, whole cell lysate (WCL) as well as cytosolic (C), nuclear (N), and crude mitochondrial (M) fractions were collected for immunoblot analysis.

### Topology analysis of mitochondrial proteins

Alkaline carbonate extractions at pH 10, pH 11 or pH 12 from crude mitochondria were performed as previously described (Wettmarshausen *et al*, 2018) to analyse membrane association and sub-mitochondrial localization of proteins. Briefly, 100 µg of mitochondria were centrifuged at 8000x*g* for 10 min at 4°C and then resuspended in 0.1 M Na_2_CO_3_ at pH of 10, 11, or 12 and incubated for 30 min on ice. Afterward, the samples were centrifuged at 45000x*g* for 10 min at 4°C. Pellets resulting from this process were resuspended in 100 µL of 2x Laemmli buffer, boiled at 98°C for 5 min, and stored at –80°C for subsequent use (referred to as the membrane sample). Supernatants were combined with 40 µL of 100% trichloroacetic acid and left to incubate overnight at –20°C. The next day, the supernatants were centrifuged at 16000x*g* for 25 min at 4°C. Pellets were washed twice with cold acetone, air-dried for 20-30 min at RT, resuspended in 100 µL of 2x Laemmli buffer, and heated to 98°C for 5 min (referred to as the soluble sample). SDS-PAGE analysis was performed on 25 µL of both the soluble and membrane samples. Antibodies against MICU1 (soluble, membrane associated), ATP5a (soluble, membrane associated), MCU (integral transmembrane protein) were used as positive controls. To determine the sub-mitochondrial localization of MICU3 and EFHD1, 30 µg of crude mitochondria were exposed to increasing concentrations of digitonin or 1% Triton X-100, which sequentially permeabilize the outer and inner membranes. This was conducted in the presence of 5 mM membrane-impermeable maleimide functionalized polyethylene glycol (mPEG, Sigma-Aldrich, 63187). This compound attaches a polyethylene glycol polymer chain of approximately 5 kDa to free thiol groups of proteins. The reaction was carried out at RT for 30 min and was quenched with 100 µM cysteine on ice for 10 min. Immunoblot analysis was performed on the samples, with the intermembrane space protein MICU1, and the matrix soluble proteins HSP60 and SOD2, serving as controls. The proteinase K protection assay was performed by incubating 30 µg of mitochondria in 30 µL of isolation buffer in the presence of 100 µg/mL proteinase K with increasing concentrations of digitonin or 1% Triton X-100 to sequentially permeabilize outer and inner membranes. The reaction was carried out at RT for 15 min and was stopped by the addition of 5 mM phenylmethylsulfonylfluorid (PMSF), followed by incubation on ice for 10 min. Immunoblot analysis was used to examine the samples, with TOM20 and TIM23 (integral outer and inner membrane proteins, respectively), along with cyclophilin D (CyPD) and HSP60 (soluble matrix proteins) as controls.

### Blue native page analysis of mitochondrial protein complexes

Blue-Native (BN) PAGE analysis was performed as described by Witting et al. (Wittig *et al*, 2006) with some adaptations. Briefly, equal amounts of mitochondria (10 µg per lane, unless noted otherwise) were diluted at least 1:100 in ice-cold miliQ water with proteinase inhibitors (Sigma-Aldrich, 5056489001), the pellet was then collected at 20000x*g* for 10 min at 4°C, resuspended in ice-cold 1x BN Sample Buffer (Thermo Fisher Scientific, BN2003) with 1% (w/v) digitonin and incubated on ice for 15 min. Afterwards, the sample was centrifuged at 20000x*g* at 4°C for 30 min and the supernatant was transferred into a new pre-chilled tube with 0.25% G-250 (Thermo Fisher Scientific, BN2004). BN-PAGE was performed at 4°C on Native PAGE Novex 3-12% Bis-Tris Protein gels (Thermo Fisher Scientific, BN1001) according to manufacturer’s instructions, followed by overnight wet blot transfer at 30 V and 4°C onto a 0.2 µM pore size PVDF membrane (Amersham, GE10600021). Immunoblotting was performed according to the standard protocol. Second dimension (2D) analysis was performed as described by Na Ayutthaya et al. (Na Ayutthaya *et al*, 2020) with some modifications. Following the above described first dimension (1D) BN-PAGE, sample lanes for 2D were excised, incubated in 1x SDS sample buffer (Bio-Rad, 1610747) for 10 min and boiled shortly, followed by incubation in hot 1x SDS sample buffer for 15 min. As a control, one well was loaded with 5 µg of input mitochondria previously diluted in 5 µL of 2x LDS dye (Thermo Fisher Scientific, 84788) with 2.75 mM β-mercaptoethanol (Thermo Fisher Scientific, 21985-023) and RIPA buffer (total volume 10 µL) and boiled for 5 min at 95°C. The 2D well of the pre-cast NuPAGE 4%-12% Bis-tris 2D well gel (Thermo Fisher Scientific, NP0326BOX) was washed and filled with 1x MOPS running buffer (Thermo Fisher Scientific, NP0001). Lanes were then fitted onto the 2D and overlaid with 1x SDS sample buffer, followed by electrophoresis according to manufacturer’s instructions. The proteins were then transferred by wet blot transfer on 0.2 µM pore size PVDF membrane (Amersham, GE10600021) and immunoblotted using MCU (Sigma-Aldrich, HPA016480) according to the standard protocol.

### RNA extraction and quantitative real-time PCR

RNA was isolated with the RNeasy Mini kit (Qiagen, 74104), according to manufacturer instructions. An equal amount of RNA from each sample was used to generate complementary DNA (cDNA) with the SuperScript™ III First-Strand Synthesis SuperMix kit (Thermo Fisher Scientific, 18080400). The resulting cDNA was diluted 1:8 in nuclease-free water and analyzed by real-time PCR using the following TaqMan assays: Hs00368816 (EFHD1) and Hs01003267 (HPRT1, used as control).

### Mitochondrial bioenergetics

HeLa sh5-EFHD1 and pLKO mt-AEQ cells were seeded at a density of 20000 cells/well in a Seahorse XFe96 Cell Culture Microplate 24 h before the experiment in DMEM with 10% FBS. On the day of the experiment, medium was removed, and cells were washed twice with 200 µL of the respective assay medium for mito-stress test (Agilent Base Medium supplemented with 1mM pyruvate, 2mM L-glutamine, 10mM glucose, pH 7.4) or glycolysis (Agilent Base Medium supplemented with 2mM L-glutamine, pH 7.4). Next, 180 µL of the assay medium for mito-stress test or glycolysis were added to each well and the plate was incubated for 60 min in a non-CO_2_ incubator at 37°C. For the mito-stress test, 10x port solution of the following compounds were injected to reach the specified final concentration: oligomycin (1.5 µM), carbonyl cyanide m-chlorophenyl hydrazone (CCCP; 1.5 µM), and antimycin A/rotenone (4 µM/2 µM). For the glycolysis assay, glucose (10 mM), oligomycin (1.5 µM), and 2-DG (100 mM). The CyQUANT® Cell Proliferation Assay Kit (Thermo Fisher Scientific, C7026) was used to normalize differences in cell density between wells. Briefly, growth media was removed, and the plate was frozen at –80°C. Cells were thawed and lysed with 200 µL of dye/lysis buffer. After 5 minutes, fluorescence was measured at 480ex/520em using a CLARIOstar plate reader (BMG Labtech). The raw fluorescence of each well was divided by the plate average and the resulting factor was used to normalize the Seahorse data.

### Cell Viability

12500 cells/well from HeLa pLKO and HeLa sh5-EFHD1 were seeded into 96-well plates. Resazurin sodium salt (Santa Cruz Biotechnology, sc-206037) was used to assess cell viability. For untreated cells, culture media was changed 48 h after seeding to 0.004% resazurin in DMEM 10% FBS medium and incubated for 4 h at 37°C after which the fluorescence was measured at 540ex/590em using a CLARIOstar Plate Reader (BMG Labtech). For treatments, 24 h after seeding, both cell lines were treated with 0.5 mM H_2_O_2_ (Sigma Aldrich, 516813), 500 nM thapsigargin (Sigma Aldrich, 586005), 40 µM C2-Ceramide (Santa Cruz Biotechnology, sc-201375) or 50 nM Paclitaxel (Abcam, ab120143-10mg), and 48 h after treatment cell viability was assessed as described above.

### Cancer Data Sources

RNA-Seq and proteomics data were obtained from the DepMap portal (https://depmap.org/), RNA-Seq gene expression TPM values from DepMap Public 23Q2 release (Ghandi *et al*, 2019), and proteomics from the Nusinow et al., 2020 data release (Nusinow *et al*, 2020). Primary breast tumor and adjacent normal tissue gene expression pair data was analyzed with TNMplot.com (Bartha & Győrffy, 2021) using the RNA-Seq data as source. Kaplan-Meier plot to assess overall breast cancer patient survival was generated using GEPIA2 (Tang *et al*, 2019), selecting the BRCA dataset for analysis and setting the median expression value as cutoff for determining the high and low expressing EFHD1 cohorts. Gene expression data for chemotherapy responders and non-responders was analyzed from rocplot.org (Fekete & Győrffy, 2019) selecting the pathological complete response cohort, any chemotherapy, and JetSet only data as parameters.

### Cell Migration Assay (Boyden Chamber Assay)

Migration assays were performed using 24-well plates with uncoated polycarbonate membrane inserts (BD Biosciences, 353097). 2×10^5^ cells were allowed to migrate for 24 h. Membranes were fixed with Methanol (Merck, 106009) stained by Hematoxylin solution (Sigma, 51275) and mounted on glass slides with Pertex mounting medium (#41-4010-00, MEDITE GmbH). Images were taken with an Olympus BX43 microscope.

## ACKNOWLEDGEMENTS

We thank Tullio Pozzan for his substantial contribution to this study, both in the design of the experiments and interpretation of the results, until his passing in October 2022. We thank Marcus Conrad, Toshitaka Nakamura, Suresh Joseph and David Weaver for critical reading of the paper and helpful discussions; Tim König and Daniela M. Arduino for experimental advice. F.P., H.C.D., S.W., N.P.D., M.F., Y.C., and M.C. were supported by the Munich Center for Systems Neurology (SyNergy EXC 2145; Project ID 390857198) and the ExNet-0041-Phase2-3 (‘SyNergy-HMGU’) through the Initiative and Network Fund of the Helmholtz Association. D.V.R. was supported by European Union funding program Horizon Europe (HORIZON-MSCA-2021-PF Project ID 101065790). J.W., A.L., and M.G. were supported by the German Research Foundation (DFG) under the Emmy Noether Programme (PE 2053/1-1) and the Bavarian Ministry of Sciences, Research and the Arts in the framework of the Bavarian Molecular Biosystems Research Network (D2–F5121.2–10c/4822) M.K. and G.H. were supported by an NIH grant (RO1-HL142271) to G.H. The authors acknowledge Euro-BioImaging (www.eurobioimaging.eu) for providing access to imaging technologies and services via the ALM Node (Padua, Italy).

## AUTHOR CONTRIBUTIONS

Conceptualization, M.M., F.P.; Methodology, M.M., F.P., G.H., H.C.D., D.V.R.; Formal Analysis, M.M., F.H., Y.C.; Visualization, S.W., Y.C., H.C.D., J.W., D.V.R.; Investigation, H.C.D., D.V.R., M.K., J.W., A.L., M.P., E.G., M.G., F.H., M.M., H.M., N.D.M., M.C., M.F. Resources, F.P., M.M., G.H., T.L., N.P., D.M.; Writing – Original Draft, F.P., H.C.D., D.V.R.; Supervision, M.M., F.P., G.H.; Funding acquisition: F.P.

## SUPPLEMENTARY TABLES

**Table S1**. List of MCUC interactors identified in MCU-TAP, EMRE-TAP, and MCUB-TAP.

**Table S2**. Protein expression level of MCUC interactors in human tissues.

**Table S3**. Protein phylogeny of MCUC interactors across Eukaryotes.

**Table S4**. List of MCU interactors upon silencing of MICU1, MICU2, EMRE, and MCUB.

**Table S5**. Pathway Enrichment Analysis of MCU interactors upon silencing of MICU1, MICU2, EMRE, and MCUB.

## SUPPLEMENTARY FIGURES

**Figure S1.** Expression of MCUC components before and after bait induction. Immunoblot analysis of MCU, EMRE, MCUB, MICU1, MICU2 and ACTIN (loading control) in whole cell lysates from Flp-In T-REx HEK293 cell lines before (–Tet) and after (+Tet) tetracycline-driven expression of each bait. Refer to quantification in Fig. 2A.

**Figure S2.** MICU3 positively regulates MCU-dependent mitochondrial Ca^2+^ uptake and PRELID1 is required for MCUC stability. **A**, **B**, MICU3 is enriched in mitochondria from (A) HeLa cells overexpressing MICU3-V5 and (B) mouse brain. *WC*, whole cell lysate; *C*, cytosol; *N*, nuclei; *M*, mitochondria. MICU1, ATP5A, MCU, and VDAC are used as markers of mitochondrial proteins, while ACTIN and LAMIN as markers of cytosolic and nuclear proteins, respectively. **C**, Immunoblot analysis of mitochondria isolated from mouse brain in presence of increasing concentrations of the membrane impermeable sulfhydryl group reactive PEG derivative (mPEG, maleimide functionalized polyethylene glycol). MICU1 is used as positive control for IMS proteins, whereas HSP60 and SOD2 for mitochondrial matrix proteins. **D**, Immunoblot analysis of mitochondrial soluble (S) and membrane (M) fractions isolated from HeLa cells overexpressing MICU3-V5 by alkaline carbonate extraction at pH 10, pH 11, and pH 12. MICU1 and ATP5A (soluble and membrane associated proteins, respectively), and MCU (integral transmembrane protein) are used as positive controls. **E, F**, MICU3 and MICU1 dimerize through a disulfide bond in mitochondria of (E) HeLa cells overexpressing MICU3-V5 compared to control (MOCK) and of (F) mouse brain. Immunoblot analysis was performed in both reducing (+DTT) and non-reducing (–DTT) conditions. *Indicates non-specific bands. MICU1/MICU3 dimers are indicated by an arrow. **G**, Domain structure of MICU3. EF1 and EF2 refer to two evolutionarily conserved EF-hand domains. Amino acid substitution used to generate MICU3 EF-hand mutants are indicated in red (EF1_mut_, D245A and E256K; EF2_mut_, D483A and E494K). **H**, Immunoblot analysis of exogenous MICU3 detected with an anti-V5 antibody and ACTIN (loading control) in whole cell lysates from HeLa mt-AEQ cells expressing either WT MICU3 (MICU3) or MICU3 mutants in the first (MICU3_EF1_), the second (MICU3_EF2_) or both (MICU3_EF1+2_) EF-hands fused to a C-terminal V5 tag and compared to untransfected control cells (MOCK). **I**, Average traces and quantification of [Ca^2+^]_mt_ transients in HeLa mt-AEQ cells expressing either WT or MICU3 mutants in response to histamine (Hist) and compared to control (MOCK). Data represent mean ± SEM (n= 4); one-way ANOVA with Dunnett’s multiple comparison test (***p< 0.001; *ns*, not significant). **J**, Immunoblot analysis of MCU, MICU1 and MAIP1 (loading control) in whole cell lysate from HeLa cells transfected with si-PRELID1 and compared to negative control (Scr). si-MCU and si-EMRE are used as positive controls for MCUC expression and stability. **K**, Representative traces and quantification of [Ca^2+^]_cyt_ transients upon histamine (Hist) stimulation in si-PRELID1 and Scr HeLa cells expressing cytosolic aequorin (mean ± SEM; n= 3); student’s t-test (*ns,* not significant). **L**, JC1-based quantification of mitochondrial membrane potential upon PRELID1 knockdown in HeLa cells (RFU, relative fluorescence unit), (mean ± SEM; n= 3); student’s t-test (*ns,* not significant). Refer to Fig. 5.

**Figure S3.** EFHD1 inhibits MCU-dependent uptake of Ca^2+^ in mitochondria without affecting [Ca^2+^]_cyt_ transients. **A**, Quantification of EFHD1 KD by real-time PCR (mean ± SEM; n= 6); one-way ANOVA with Dunnett’s multiple comparisons test (****p< 0.0001). **B**, **C**, Representative traces and quantification of [Ca^2+^]_cyt_ transients upon histamine (Hist) stimulation in (B) sh-EFHD1 and (C) si-EFHD1 HeLa cells expressing cytosolic aequorin (mean ± SEM; n≥ 12); one-way ANOVA with Dunnett’s multiple comparisons test (*ns*, not significant). **D**, Quantification of [Ca^2+^]_mt_ (upper panel) and [Ca^2+^]_cyt_ (lower panel) responses in control (Scr) and si-EFHD1 treated HeLa cells upon histamine-induced ER Ca^2+^ release in presence of 1.8 mM Ca^2+^ in the extracellular medium (ECM). Peak and area under the curve (AUC) are calculated for the first 60 s of histamine (Hist) stimulation (mean ± SEM from 3 independent experiments (n≥ 30)); one-way ANOVA with Dunnett’s multiple comparisons test (***p< 0.001, ****p< 0.0001; *ns*, not significant). Refer to Fig. 4.

**Figure S4.** Effect of EFHD1 knockdown on MCUC protein stability and assembly. **A**, Immunoblot analysis of MCUC protein level in whole cell lysates from HeLa cells upon stable (sh-EFHD1) or transient (si-EFHD1) EFHD1 silencing. GRP75 is used as a loading control. **B**, BN-PAGE analysis of MCUC assembly in mitochondria isolated from sh-EFHD1 HeLa cells. ATP5A is used as a loading control. **C**, Immunoblot analysis of EFHD1 in isolated mitochondria from HeLa cells treated with increasing concentrations of digitonin and maleimide functionalized polyethylene glycol (mPEG). MICU1 and SOD2 are used as positive controls for IMS and matrix proteins, respectively. **D**, Immunoblot analysis of mitochondria from HeLa and HEK293 cells expressing either an empty vector (pLKO) or shRNA against EFHD1. Samples were analyzed in reducing (+DTT, dithiotreitol) and non-reducing (–DTT) conditions to detect disulfide-mediated oligomerization. Refer to Fig. 5.

**Figure S5.** Assessment of EFHD1 as a potential target in cancer. **A**, Heatmap of gene expression level (TPM, transcripts per million) for the known MCUC components and EFHD1 retrieved from the DepMap Public 23Q2 release (Ghandi *et al*, 2019) and grouped according to cell line lineage (n= 1450 cell lines; 29 lineages). Averaged values are inferred from RNA-sequencing data using the RSEM tool and log_2_ transformed, using a pseudo-count of 1 (log_2_(TPM+1)). **B**, Correlation between protein and RNA levels of EFHD1 in 1019 different cancer cell lines, grouped based on cell lineage average expression. RNA-sequencing data were retrieved from DepMap Public 23Q2 release (Ghandi *et al*, 2019) whereas normalized protein expression data were taken from (Nusinow *et al*, 2020); RNA-protein expression Pearson correlation r^2^=0.85. **C**, Median EFHD1 expression in pairs of primary breast tumor and their adjacent normal tissue (n= 112). Data were extracted from TNMplot.com median ± 95% confidence interval (CI) and Wilcoxon match-paired two-tailed test; ****p< 0.0001. **D**, Median EFHD1 expression in neoadjuvant chemotherapy in responder (n= 532) and non-responder (n= 1100) breast cancer patients analyzed using ROCplot.org from GEO/Array express data median ± 95% CI; Mann-Whitney two-tailed test; *p< 0.05. **E**, Survival of breast cancer patients exhibiting high and low EFHD1 expression. Data were retrieved from TCGA-BRCA; Kaplan-Meier plot and the median expression was used as the cohort cut-off, respectively. *HR*, hazard ratio. Dotted lines indicate 95% confidence interval (CI 95%). **F**, Boyden Chamber migration assay on EFM19 pLKO and sh-EFHD1 cells. Representative image (left) and quantification of migrated cells per field (right). Mean ± SEM from 3 independent experiments (n≥ 30); student’s t test; ns, not significant. Refer to Fig. 5.

**Figure S6.** Validation of MCUB as an inhibitor of MCU protein-protein interactions. **A**, Immunoblot analysis of MCUB in whole tissue lysates from MCUB KO and wild-type mice. Blue arrow indicates MCUB protein. **B** Relative MCUB protein expression in mouse tissues from the ProteomicsDB database (Normalized iBAQ). **C**, Relative MCUB protein expression in pure mitochondrial proteomes extracted from Hansen et al., 2023. MCUB was detected only in spleen and brain. Refer to Fig. 6.

## Notes

### Competing Interest Statement

The authors have declared no competing interest.

