## Supplementary figures and images for "Systematic mapping of MCU-mediated mitochondrial calcium signaling networks"

### Supplemental Figure 1

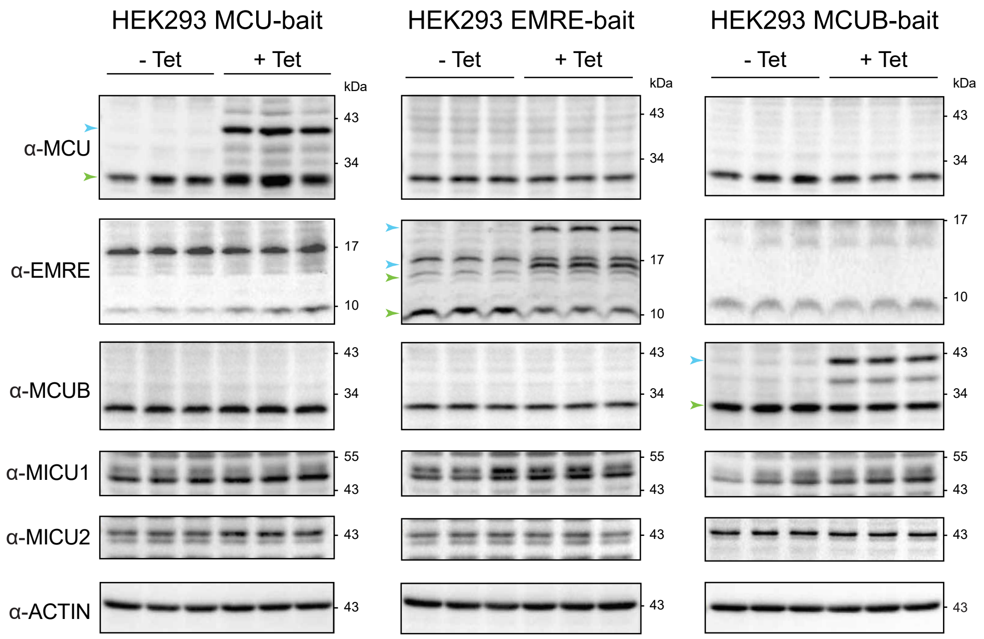


Delgado et al._Figure S1

### Supplemental Figure 2

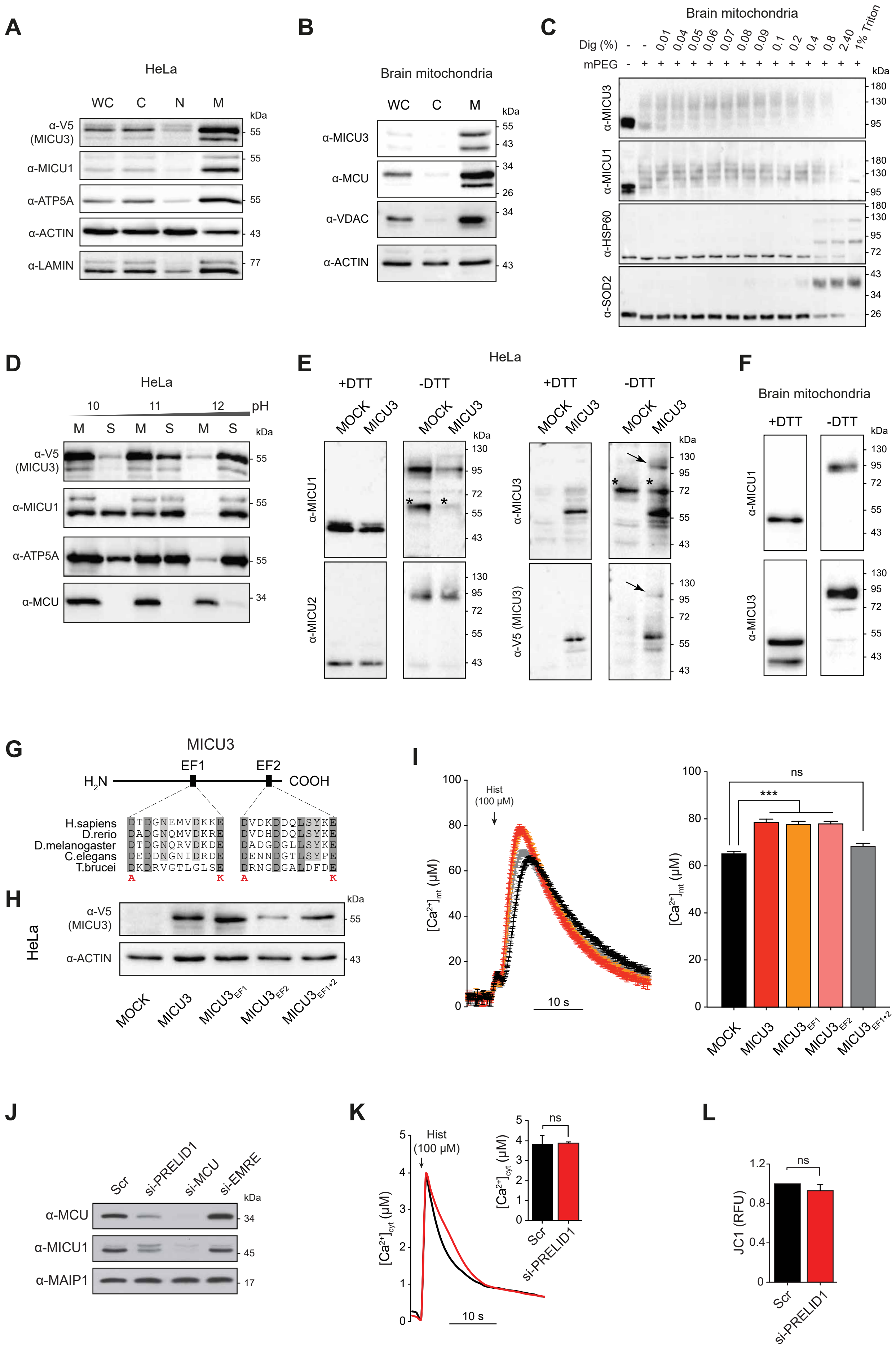

Delgado et al. Figure S2

### Supplemental Figure 3

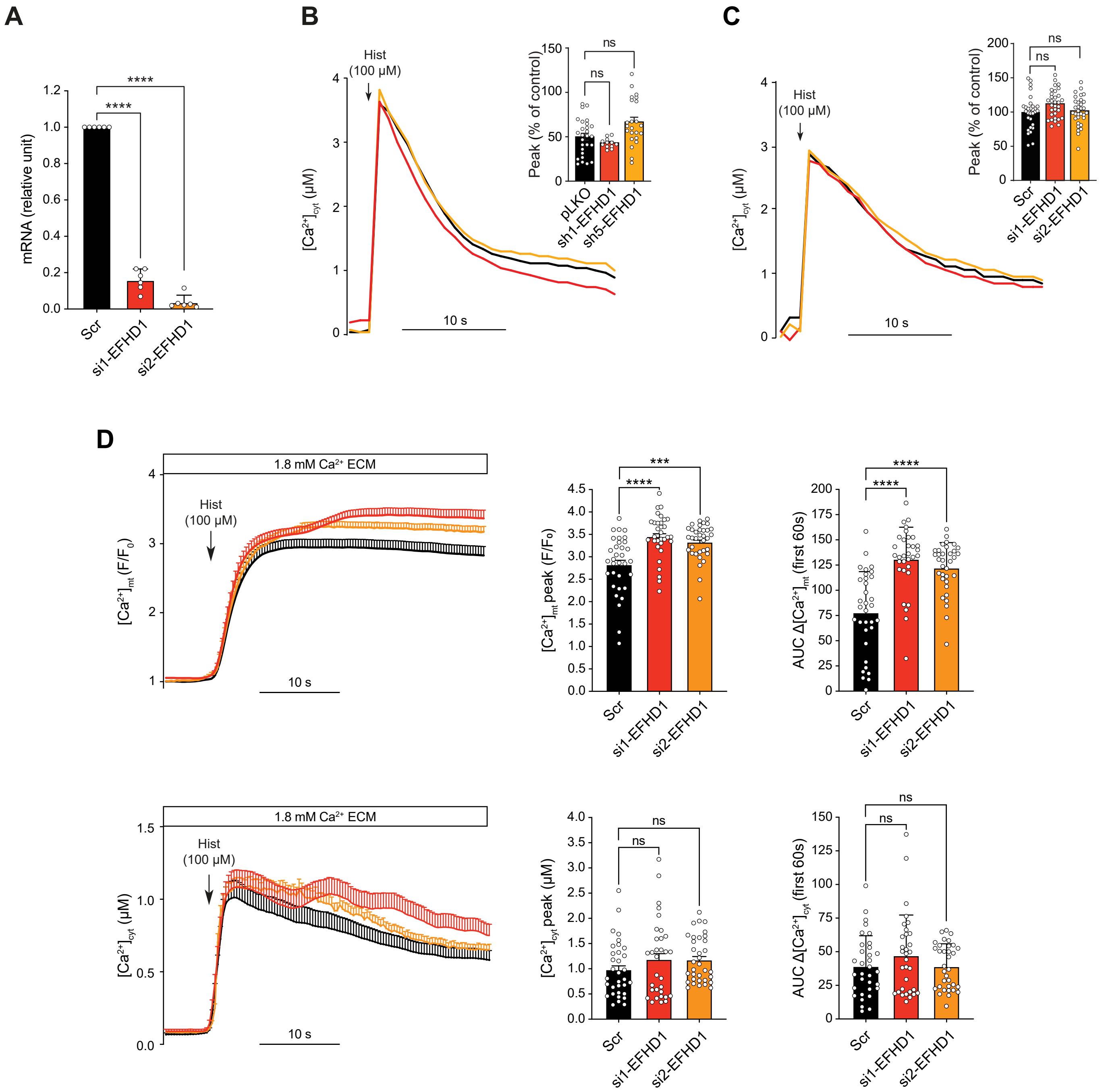


Delgado et al._Figure S3

### Supplemental Figure 4

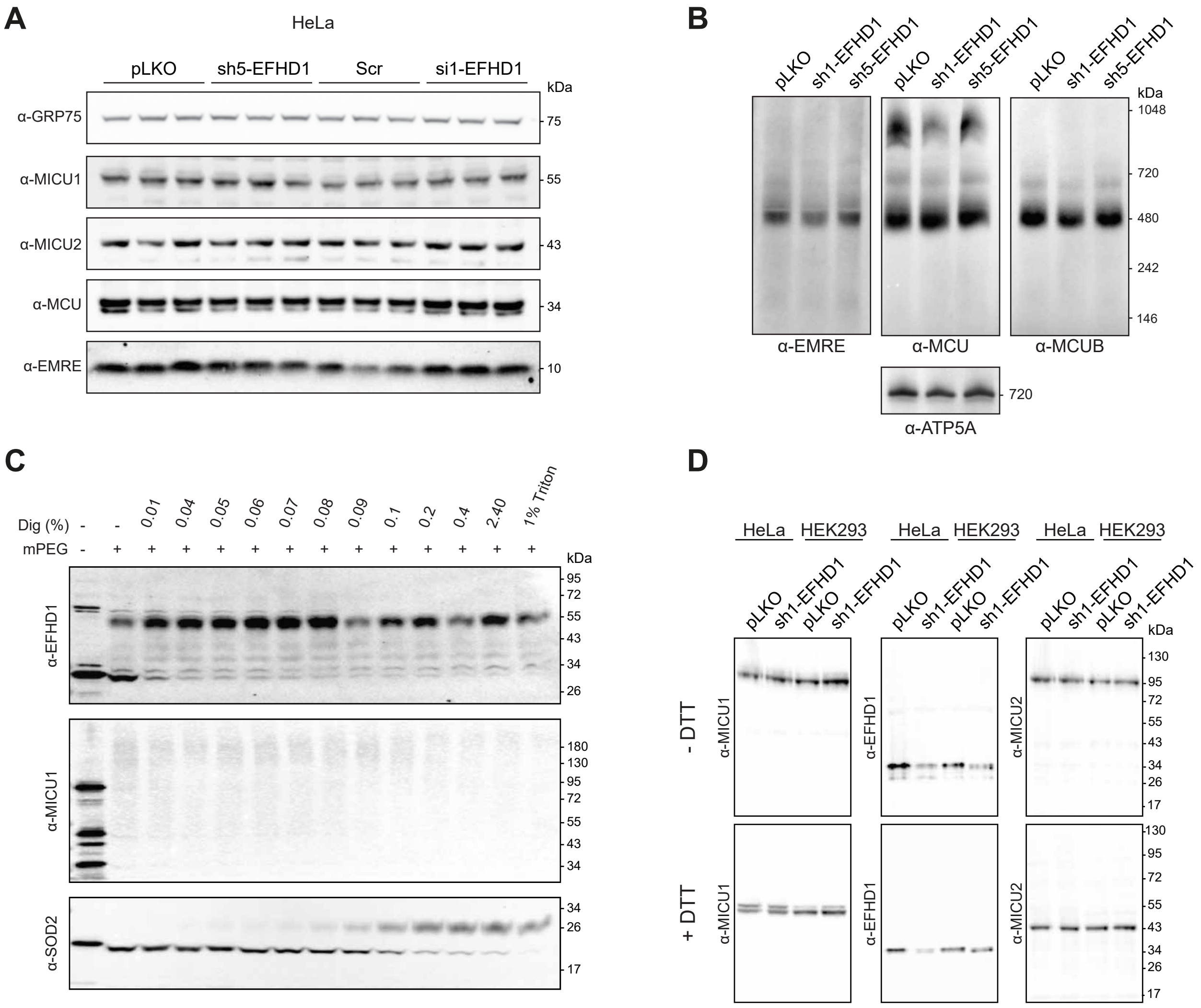


Delgado et al._Figure S4

### Supplemental Figure 5

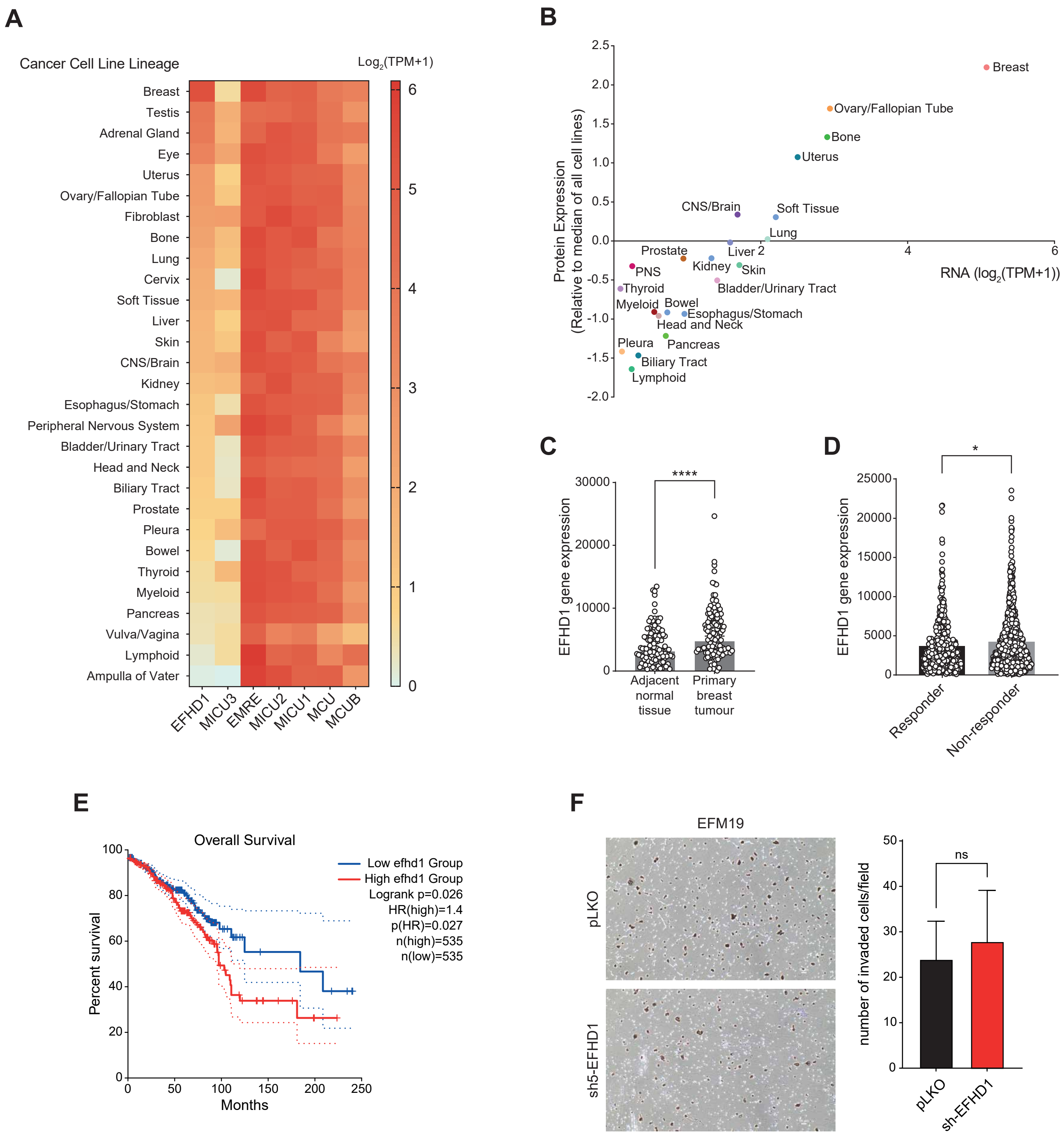


Delgado et al. Figure S5

### Supplemental Figure 6

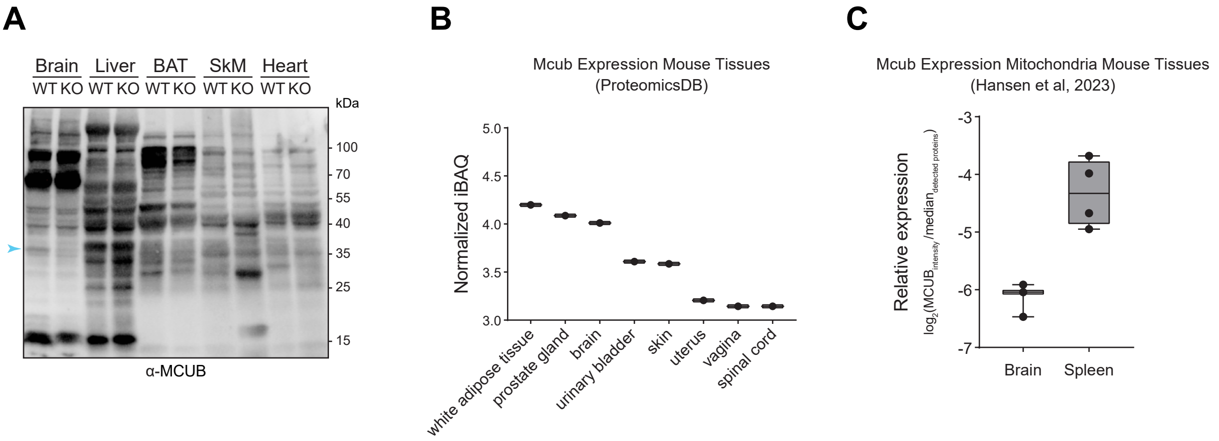


Delgado et al._Figure S6
